# A data-driven approach to identifying and evaluating connectivity-based neural correlates of conscious visual perception

**DOI:** 10.1101/2025.04.06.646695

**Authors:** Annie G. Bryant, Christopher J. Whyte

## Abstract

Identifying the neural correlates of conscious visual perception remains a major challenge in neuroscience, requiring theories that bridge between subjective experience and measurable neural correlates. However, theoretical interpretation of empirical evidence is often post hoc and susceptible to confirmation bias. Building upon the adversarial collaboration mediated by the COGITATE Consortium, we present a generalizable approach for the data-driven identification, evaluation, and theoretical modeling of connectivity-based neural correlates of conscious visual perception. Using the same magnetoencephalography (MEG) dataset and accompanying pre-registered hypotheses from the COGITATE Consortium, we systematically compared 246 functional connectivity (FC) measures between regions predicted to underlie conscious vision by Integrated Information Theory (IIT) and/or Global Neuronal Workspace Theory (GNWT). We identified a family of FC measures based on the barycenter—tracking the ‘center of mass’ between two signals—as the top-performing stimulus decoding measures that generalize across regions central to predictions of both IIT and GNWT. To interpret these findings within a theoretical framework, we developed neural mass models that recapitulate the neural dynamics hypothesized to underlie conscious perception by each theory. Comparing simulated barycenter values from these models against empirically measured MEG data revealed that both the GNWT-based model, featuring delayed ignition dynamics, and the IIT-based model, which relied on synchronous sensory dynamics, captured the observed connectivity patterns. These results lend tentative support to GNWT, as the presence of ignition dynamics independent of task-demand conditions contradicts the predictions of IIT. Beyond dataset-specific conclusions and limitations, we introduce a framework for systematically identifying and testing candidate neural correlates of conscious visual perception in an unbiased and interpretable manner.

## 1 Introduction

Characterizing the neural correlates of conscious visual perception remains one of the great open challenges in modern neuroscience. At the heart of this challenge is the search for a theory of consciousness that can map between subjective experiences and their observable neural correlates [1]. Although the field sees an increasing number of publications each year, theoretical interpretation is primarily performed post hoc; moreover, design choices of the methodology itself are highly predictive of which theory the study’s findings will support [2]. These biases have cultivated a fragmented literature that lacks a cohesive methodology, in terms of both experimental paradigms and theoretical interpretation.

In a collective endeavor to move beyond current theoretical silos and confirmation biases, researchers from competing theoretical camps have jointly entered into adversarial collaborations that directly test their preregistered predictions in shared empirical paradigms, conducted by theory-neutral third-party experimental groups. This adversarial collaboration, led by the COGITATE Consortium, focused on the predictions of two prominent theories of consciousness: integrated information theory (IIT) and global neuronal workspace theory (GNWT). IIT identifies consciousness with the emergent cause–effect structure of physical systems [3]. According to IIT, a system is conscious to the degree that the system as a whole exhibits greater informational content than the sum of information contained within all sub-partitions of the system. In contrast, GNWT [4, 5] identifies consciousness with the availability of information in the ‘global workspace’ [6], a recurrent network with long-range excitatory projections connecting the prefrontal cortex (PFC) and parietal cortices.

Within the domain of conscious vision in particular, IIT and GNWT can be cast as yielding contrasting predictions regarding stimulus decodability, the location and temporal dynamics of conscious vision, the role of pre-stimulus activity, and the nature of inter-areal coupling. Advocates of IIT involved in the COGITATE Consortium postulate that consciousness is a persistent structure, with conscious perception requiring phase synchronization between low-level visual areas (including primary visual cortex) and category-selective visual areas (lateral occipital cortex [7] and fusiform face area [8]). Together with regions in the parietal cortex, this network is often referred to as the ‘posterior hot zone’—and notably does not include the PFC, which IIT advocates argue is only necessary for cognitive aspects of consciousness, such as introspection and self-report [9]. By contrast, advocates of GNWT involved in the COGITATE Consortium posit that stimulus information becomes conscious when it enters the global workspace through a late wave of neural signal propagation from sensory cortices to PFC. This wave comprises a nonlinear ignition event, independent of task demands, through which the global workspace broadcasts information between otherwise isolated processes throughout the brain. In the experiments conducted by the COGITATE Consortium [10], the results of which were recently made available [11], these predictions were tested through a shared visual perception paradigm agreed upon by theorists from both IIT and GNWT. Specifically, participants were shown a stream of images belonging to one of four categories (face, object, letter, or false font). Across multiple functional neuroimaging modalities, the COGITATE Consortium reported varying degrees of evidence supporting the predictions put forward by GNWT and IIT theorists based on region-specific decoding and the timing of conscious percepts. However, neither GNWT nor IIT theorists’ predictions about the relationship between inter-areal coupling (assessed via functional connectivity) and stimulus decodability were borne out—raising the question of whether there exists a functional connectivity (FC) measure that can discriminate between the neural correlates of conscious visual experience predicted by IIT and/or GNWT.

Of note, the COGITATE Consortium only examined two measures of inter-areal connectivity: pairwise phase consistency [12] and dynamic FC ([13]; using the Pearson correlation coefficient) between respective pairs of theory-predicted brain regions. Many other measures of inter-areal coupling could, in principle, distinguish between the predictions of each theory in the relevant region–region pairs. For example, in previous work, we systematically compared diverse coupling types—for example, synchronous vs. asynchronous, directed vs. undirected, and time vs. frequency domain-based [14, 15]. The results validated commonly used measures like the Pearson correlation coefficient across neuroimaging modalities, while simultaneously highlighting the potential utility of previously unexplored types of measures of pairwise interaction, such as directed information [16]. Moreover, the predictions put forward by both IIT and GNWT theorists were intrinsically qualitative, making it difficult to precisely quantify the evidence for predictions from either theoretical camp.

In this work, we address these limitations in the quantitative testing of competing theories by presenting a unified and data-driven approach for identifying, evaluating, and theoretically interpreting candidate connectivity-based measures of the neural correlates of conscious visual perception. Based on the aims of the COGITATE Consortium [11] pertaining to inter-areal coupling, we present a highly comparative approach to search for inter-areal coupling (i.e., FC) measures that reliably quantify the neural correlates of conscious vision from magnetoencephalography (MEG) imaging data. This comprehensive suite of 246 pairwise coupling measures [14] includes those typically employed in functional neuroimaging, including the Pearson correlation coefficient, pairwise phase consistency, power envelop correlation [17, 18], and *ϕ*^∗^ [19]—enabling direct comparison to under-explored alternatives. While it is unlikely that a single FC measure is the optimal choice across all comparisons in a dataset, this framework provides a meta-comparative approach to identify broad classes of measures that perform well across a variety of tasks, which can in turn inform future theoretical applications. In particular, we found high overall performance of FC measures derived from the barycenter [20], a center-of-mass statistic originally developed in astrophysics that is sensitive to both amplitude- and phase-based coupling between two brain regions. The barycenter is computed as a new time series that minimizes the distance to the MEG activity in two given regions using a particular distance metric, which can include Euclidean and non-Euclidean geometries such as dynamic time warping. The resulting barycenter can be used to derive quantities such as the maximum squared magnitude, reflecting the maximal covariance in MEG activity (according to a given distance metric) of the two regions. To our knowledge, this is the first application of barycenter-derived FC for stimulus decoding in the human brain, underscoring the benefit of our data-driven framework to highlight novel biologically relevant properties of neural activity.

We extend this (preliminary) finding of a generally informative FC measure family in distinguishing visual stimuli to quantitatively interpret the FC results within the theoretical frameworks of IIT and GNWT, respectively. This is achieved using two neural mass models [21] that implement the stimulus-evoked dynamics hypothesized by each theory to be necessary for conscious perception, which simulate MEG-like regional dynamics. Crucially, implementing the hypothesized dynamics for each brain region in computational models allows us to explicitly compute exemplar time series and quantitatively compare the predictions of each theory based on the top-performing FC measures. In sum, we present a data-driven approach for identifying the FC measures that maximally encode stimulus-specific information, and then implement simple and interpretable neural models to derive theoretical insight from these FC measures.

## 2 Materials and methods

### 2.1 Neuroimaging data acquisition and task paradigm

All data was obtained from the COGITATE Consortium [11] as prepared for a functional connectivity data science challenge (cf. https://www.arc-cogitate.com/biomag-2024) [22]. MEG data were downloaded in BIDS format for 100 participants, N=94 of whom had the requisite anatomical scans to perform source localization. This subset of N=94 participants included N=54 females (23.1 *±* 3.5 years) and N=40 males (22.3 *±* 3.6 years) with no known neurological or psychological conditions and no age differences between males and females (Wilcox rank-sum test, *P*=0.19). All imaging acquisition and behavioral task details for this dataset are provided in COGITATE Consortium et al. [11]. Briefly, participants viewed supra-threshold visual stimuli comprised of faces, objects, letters, or false fonts, for variable durations (0.5 s, 1.0 s, or 1.5 s) at different rotation angles. Participants were instructed to press a button when either of two specific stimuli were presented, such that stimulus events were classified as ‘Relevant target’ (i.e., the correct stimulus to respond to), ‘Relevant non-target’ (i.e., all other stimuli belonging to the same category as one of the target stimuli), or ‘Irrelevant’ (i.e., a stimulus belonging to a different category to both target stimuli). We included all stimulus types in our analysis, though we excluded the ‘Relevant target’ setting due to potential motor confounds. Additionally, for simplicity, we focused exclusively on the 1.0 s (‘intermediate’) stimulus duration to compare neural dynamics during one second of stimulus presentation (‘onset’) and the subsequent post-stimulus second (‘offset’). Participants were conscious of the presented stimuli in both task-relevant and irrelevant trials, and they demonstrated high hit rates and low false alarm rates, eliminating behavioral accuracy as a potential confound [11].

### 2.2 Neuroimaging preprocessing

#### 2.2.1 Anatomical preprocessing

High-resolution T1-weighted structural volumes were preprocessed with the same methods and accompanying code outlined in COGITATE Consortium et al. [11]. Briefly, cortical surfaces were reconstructed using FreeSurfer’s `recon-all` pipeline (cf. https://surfer.nmr.mgh.harvard.edu/). Scalp surfaces were also reconstructed and combined with an inner skull surface to create a single-shell boundary elements model (using the `MNE` module in Python [23]), from which the solution was used to create a volumetric forward model spanning the full brain volume with a 5 mm grid.

The cortex was parcellated by resampling the 100-parcel atlas from Schaefer et al. [24] (mapping to the 7-Network atlas from Yeo et al. [25]) to each participant’s reconstructed cortical surface using the `mri_surf2surf` function from FreeSurfer. From this atlas, we selected a parsimonious set of four brain regions of interest (per hemisphere) to represent hypotheses of conscious visual perception from IIT and/or GNWT. Briefly, GNWT predicts that conscious perception requires an initial ‘ignition’ event in high-level visual areas (i.e., CS) that is propagated to the global workspace in the PFC. By contrast, IIT predicts that conscious contents are maintained through a persistent structure between CS and low-level visual cortices (i.e., VIS). As shown in Fig. 1A and listed in Table 1, the four regions we selected include category-selective visual areas (CS; common to both IIT and GNWT), early visual areas (VIS, comprised of V1 and V2; exclusive to IIT), and lateral prefrontal cortex (PFC, comprised of Brodmann areas 9 and 46; exclusive to GNWT). Due to the combinatorial explosion that can arise from a highly comparative analysis, we included an additional parietal region (PAR, comprised of the intraparietal sulcus), as the parietal cortex plays an important role in consciousness in both theories. Specifically, the parietal cortex is a part of the posterior substrate of consciousness according to IIT [3], and it is also a part of the ‘global workspace’ according to GNWT [5]. By including PAR as an additional region, we gained a further constraint on high-performing FC measures without biasing our search toward either theory.

**Table 1.** The individual brain regions from the 100-parcellation atlas from Schaefer et al. [24] comprising our regions of interest. These regions are highlighted on the cortical surface in Fig. 1A.

|  | Left hemisphere | Right hemisphere |
| --- | --- | --- |
| **Category-Selective (CS)** | 7Networks_LH_DorsAttn_Post_1-lh | 7Networks_RH_Vis_3-rh |
| **Primary visual (VIS)** | 7Networks_LH_Vis_4-lh | 7Networks_RH_Vis_4-rh |
|  | 7Networks_LH_Vis_5-lh | 7Networks_RH_Vis_5-rh |
| **Prefrontal Cortex (PFC)** | 7Networks_LH_SalVentAttn_PFCI_1-lh | 7Networks_RH_Cont_PFCI_3-rh |
| **Parietal integration (PAR)** | 7Networks_LH_DorsAttn_Post_3-lh | 7Networks_RH_DorsAttn_Post_4-rh |

**Figure 1.**
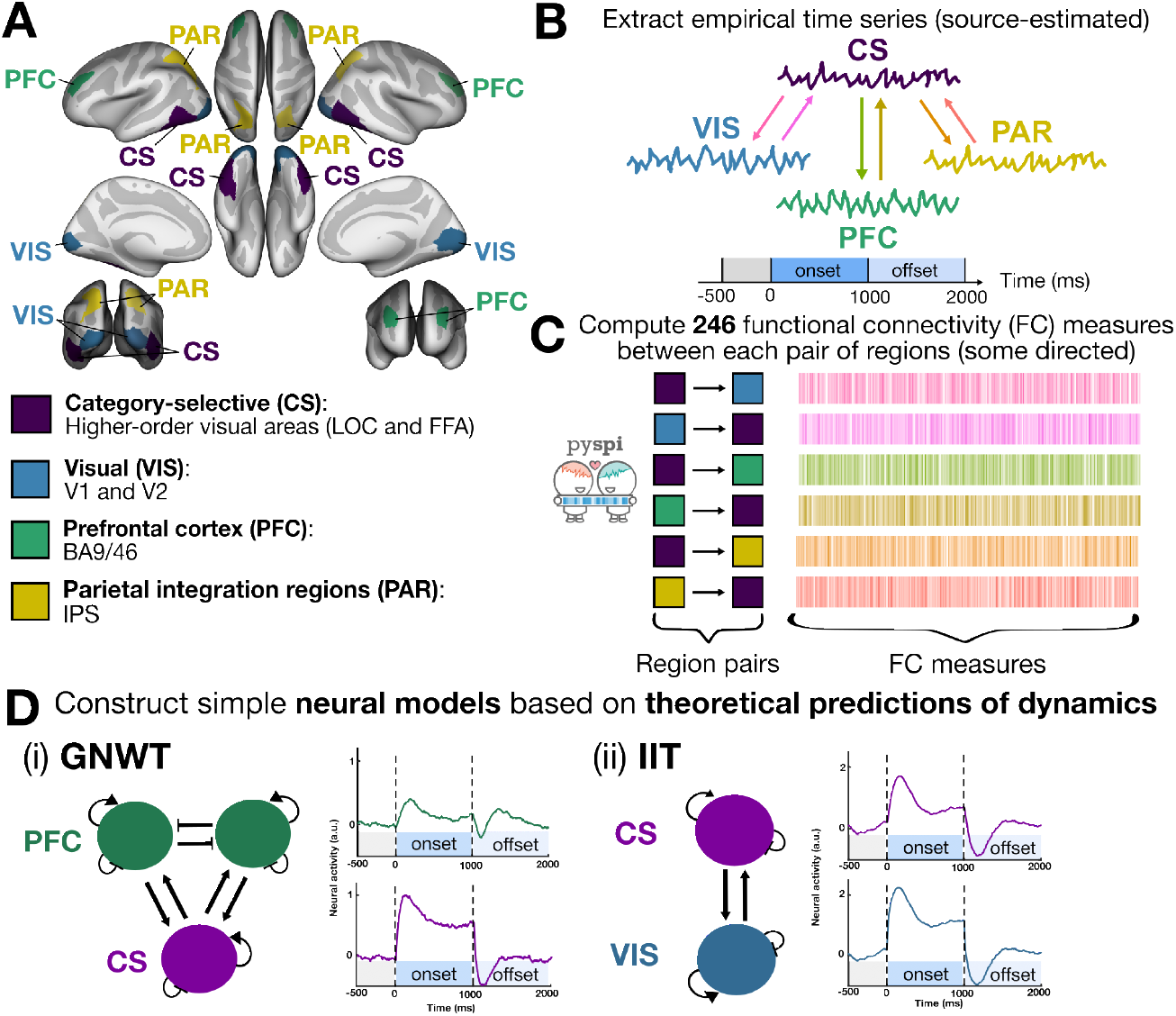
In this study, we systematically compared different ways of quantifying functional coupling from MEG time series derived from regions hypothesized to support the IIT and/or GNWT theories of visual perception and consciousness. **A**. We selected four regions of interest that are depicted on the inflated cortical surface in `fsaverage` space: Category-selective (CS), comprised of the higher-order visual lateral occipital cortex (LOC) and fusiform face area (FFA) regions; Visual (VIS), comprised of V1 and V2; Prefrontal cortex (PFC), comprised of Brodmann areas 9 and 46; and Parietal integration regions (PAR), comprised of the intraparietal sulcus (IPS). **B**. We extracted the average source-localized MEG time series across all epochs per event-related field type in each region, considering both stimulus onset (first 1000ms) and the subsequent offset period (second 1000ms). **C**. For each functional pathway (i.e., each directed region–region pair), we computed 246 statistics of pairwise interactions (SPIs) using the *pyspi* library. **D**. Neural mass models designed to recapitulate the dynamics of each theoretically relevant region based upon the preregistered predictions of GNWT and IIT set out in COGITATE Consortium et al. [11]. The dynamics of the GNWT-based model are governed by two key factors: (1) winner-take-all competition between the two PFC neural masses leading to transient ‘ignition’ events at stimulus onset and offset; and (2) PFC masses have a slower intrinsic timescale leading to the delayed peak in PFC activity. The dynamics of the IIT-based model are likewise governed by two key factors: (1) strong recurrent connectivity between CS and VIS masses; and (2) similar intrinsic timescales, which together yield highly synchronous activity at stimulus onset that is maintained for the duration of the stimulus presentation period, which then rapidly and synchronously decays at stimulus offset.

#### 2.2.2 MEG preprocessing

MEG data were preprocessed using the same methods previously employed by the COGITATE Consortium for MEG data [11, 26], which included signal-space separation with a Maxwell filter to minimize environment artifacts [27] and FAST-ICA to remove cardiac muscle artifacts [28]. Data were partitioned into 3.5 s epochs, comprised of 1 s pre-stimulus to 2.5 s post-stimulus onset, and epochs were filtered according to gradiometer, magnetometer, and/or artifact properties as described in COGITATE Consortium et al. [11]. Source modeling was performed using the dynamic statistical parametric mapping method [29]. A common spatial filter was generated from the sum of a noise covariance matrix for the baseline period (-0.5 to 0 s) and a covariance matrix for the active time window period (0 to 2 s). The baseline covariance matrix was used to spatially pre-whiten the data to combine magnetometer and gradiometer data. For each epoch, we fit and applied an inverse solution to extract the time series from the relevant regions (cf. Methods, Sec 2.2.1), averaging across all channels within each parcel. To enhance the signal-to-noise ratio [30], we averaged across trials for each event-related field type (i.e., for each stimulus type and task relevance condition) to yield one average time series per brain region for the given trial type in each participant. Given the long presentation time (1 s) combined with the supra-threshold stimulus, behavioral accuracy was close to the ceiling [11]. It is unlikely, therefore, that trial-by-trial fluctuations played a large role in task-relevant aspects of neural dynamics (justifying our averaging-based approach to maximizing signal-to-noise ratio). This assumption would need to be relaxed if one were to adapt the pipeline to typical conscious-access paradigms where pre-stimulus fluctuations are known to play an important role in explaining behavioral variability [31]. As we focused on the 1 s stimulus duration, we compared the 1 s window of stimulus presentation (‘onset’) with the following 1 s post-stimulus period (‘offset’) for each trial type.

### 2.3 Data-driven quantification of inter-areal coupling

We sought to comprehensively quantify diverse types of functional coupling between our four regions. We employed the novel framework for highly comparative analysis of pairwise interactions, `pyspi` (v1.1.0) [14], computing a total of 262 FC measures from the MEG time series between each pair of regions (e.g., CS → PAR). As part of the standard `pyspi` workflow, all time series were z-scored along the time axis, after which linear detrending was applied with the `detrend` function from the `scipy.signals` module [32]. Some of the FC measures are undirected, such that values are symmetric in both directions between region–region pairs, whereas others are directed (and therefore asymmetric). For quality control, we filtered the 262 FC measures down to those that yielded non-zero variance and real, non-NaN values across region–region pairs for at least 90% (i.e., N=85) of participants, yielding a final set of 246 FC measures. As described in Cliff et al. [14], these FC measures comprise diverse types of pairwise interactions, from basic covariance estimates to multi-metric distances to frequency-based coupling.

### 2.4 Fitting classifiers

Based on the original predictions for inter-areal coupling delineated by COGITATE Consortium et al. [11], we defined one core classification aim and two supplementary classification aims. In our main analysis targeting the core classification aim, we evaluated whether any FC measure in `pyspi` could distinguish between stimulus types (e.g., ‘face’ vs. ‘object’) in either the ‘Relevant non-target’ and/or ‘Irrelevant’ settings, in any region–region pair and during either stimulus presentation (‘on’) or offset (‘off’). For this primary classification aim (‘stimulus type’), we fit a separate classifier per FC measure for each region–region pair, task relevance condition (Relevant non-target or Irrelevant), and stimulus presentation period (‘on’ during the first 1000 ms from stimulus onset or ‘off’ during the second 1000 ms). Across all 246 FC measures, this amounted to a total of 35,280 individual models examined in this primary aim. Our secondary analyses focused on the supplementary classification aims, in order to explore whether any FC measure could distinguish domain-independent task relevance (‘Relevant non-target’ vs. ‘Irrelevant’) and/or generalize from one relevance type to another in the same stimulus comparison. For our supplementary aims (‘task relevance’ and ‘cross-task’ classification), we still partitioned the data by region–region pair (e.g., VIS → CS) and stimulus presentation period (‘on’ or ‘off’) into separate classifiers for each FC measure. As an example, for the ‘task relevance’ problem in the CS → PFC and stimulus ‘on’ setting, we grouped the participant-wise FC measure values for all four stimulus types (‘face’, ‘object’, ‘false font’, and ‘letter’) and trained the classifier to predict ‘Relevant non-target’ versus ‘Irrelevant’ regardless of domain, cross-validating with ten stratified folds across the N=94 participants. With the ‘cross-task’ problem of ‘face’ versus ‘object’ in the CS → PFC and stimulus ‘on’ setting, as an example, the classifier was trained on all N=94 participants in one relevance setting (e.g., ‘Irrelevant’) and evaluated in the other unseen relevance setting (e.g., ‘Relevant non-target’) per FC measure.

In all cases, for each FC measure (e.g., power envelope correlation), we fit a binary logistic regression classifier using *scikit-learn* (v1.6.1) [33] to predict either the stimulus type or task relevance from the associated FC measure values (from the given region–region pair). Classification performance was assessed as the mean accuracy across the test subsets per model and is presented throughout this report as the mean cross-validated accuracy *±* 1 SD. For the stimulus pair classification (primary aim) and task relevance classification (first supplementary aim), we evaluated out-of-sample performance using stratified 10-fold cross-validation with the `StratifiedGroupKFold` function from *scikit-learn*. This allowed us to train each model on a pair of stimuli (or pair of relevance conditions) from one subset of participants and evaluate the model on that pair of stimuli or relevance conditions from another (unseen) subset of participants. For the cross-task classification (second supplementary aim), each model was trained on the given pair of stimuli (e.g., ‘face’ and ‘object’) from all N=94 participants in the ‘Relevant non-target’ context, and tested on the same pair of stimuli from all N=94 participants in the ‘Irrelevant’ context (and vice versa).

While the accuracy is only computed for the default probability threshold of 0.5 for each classifier, we additionally computed the area under the receiver operating characteristic curve (AUC) in our cross-validation approach to evaluate model performance across thresholds. Across all classification problems, the accuracy and AUC exhibited similar performance (Pearson’s *R* = 0.71), so we focus on accuracy in this study to align with the binary prediction of the stimulus type (or task relevance setting, where appropriate) [34]. As a robustness check, we also fit support vector machine (SVM) classifiers with both a linear and nonlinear radial basis function kernel for each FC measure and region–pair (setting the regularization hyperparameter *C* = 1 in both cases) and evaluated the cross-validated accuracy to confirm that similar performances were achieved with the logistic regression classifiers.

### 2.5 Theory-driven neural modeling

#### 2.5.1 Model development and implementation

To quantitatively link each theory with the empirically observed neural activity (measured with MEG), we constructed two simple neural mass models [21, 35, 36] based on the neural activity predicted for each core region (PFC, CS, and VIS) defined by the COGITATE Consortium [11]. The models were derived from the Wilson-Cowan equations [37–40] under the assumption that excitatory and inhibitory currents entering the excitatory population are balanced, and that inhibitory neurons respond rapidly and are therefore well approximated by their asymptotic value [41]. We direct interested readers to the Supplementary Methods in Sec. S1 for more details. The effective dynamics were governed by the following system of ordinary differential equations:

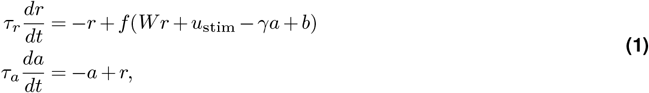

with state variables *r, a* ∈ ℝ^*N*×1^ representing the aggregate (i.e., coarse-grained) neural activity and (hyperpolarizing) adaptation current, respectively, of each cortical region hypothesized to play a role in conscious perception; for similar models applied to the visual system, we refer the interested reader to Wilson [37]. For simplicity, the transfer function *f* (*x*) is a rectified linear unit. The matrix *W* ∈ ℝ^*N ×N*^ parameterizes the strength of inter-areal connectivity with self-connections set to zero based upon the assumption of balanced intrinsic currents; the visual stimulus was encoded in the one-hot vector *u*_stim_ ∈ ℝ^*N*×1^ which, as in the experimental paradigm, lasted 1000 ms of simulation time; *τ*_*r*_ and *τ*_*a*_ are scalar time constants; *γ* is a scalar constant controlling the strength of the adaptation current; and *b* is a scalar bias parameter representing the background level of arousal.

Parameter values (supplied in Table 2) were derived from stability analysis of each model (see Supplementary Methods in Sec. S1) and were designed to recapitulate—as precisely as possible—the time series predictions for each region that were pre-registered by IIT and GNWT theorists [11] by associating each set of qualitative predictions with attractor dynamics and using stability analysis to find the parameter set over which the attractor dynamics remained qualitatively identical (i.e., where the fixed points of each dynamical system vary quantitatively but retained their stability properties) which we then swept over in simulation. Neuronal time constants (*τ*_*r*_) were selected to respect the empirical timescale [42] of the theory-relevant regions. The adaptation time constant was chosen to align with previous biophysical modeling [43]. Model simulations and analysis were implemented in custom Python (version 3.9.21) code. We solved Eq. (1) numerically using an Euler–Maruyama integration scheme (Δ*t* = 1 ms), adding zero-mean Gaussian noise (*σ* = 5.0 × 10^−3^) to both the neural activity and adaptation current.

**Table 2.** Parameter values for the simple IIT- and GNWT-based neural mass models. Parameter values are in arbitrary units unless stated otherwise. These parameters were used construct models as schematically depicted in Fig. 1D.

|  | IIT | GNWT |
| --- | --- | --- |
| $W$ | $\begin{matrix} & \text{VIS} & \text{CS} \\ \text{VIS} & \begin{pmatrix} 0 & W_{\text{IIT}} \end{pmatrix} \\ \text{CS} & \begin{pmatrix} W_{\text{IIT}} & 0 \end{pmatrix} \end{matrix}$ | $\begin{matrix} & \text{CS} & \text{PFC}_1 & \text{PFC}_2 \\ \text{CS} & \begin{pmatrix} 0 & W_{\text{GNWT}} & W_{\text{GNWT}} \end{pmatrix} \\ \text{PFC}_1 & \begin{pmatrix} W_{\text{GNWT}} & 0 & -W_P \end{pmatrix} \\ \text{PFC}_2 & \begin{pmatrix} W_{\text{GNWT}} & -W_P & 0 \end{pmatrix} \end{matrix}$ |
| $\gamma$ | 0.25 | 0.85 |
| $\tau_r$ | $\tau_{\text{VIS}} = 20, \tau_{\text{CS}} = 40$ (ms) | $\tau_{\text{CS}} = 40, \tau_{\text{PFC}} = 80$ (ms) |
| $\tau_a$ | 500 (ms) | 500 (ms) |
| $b$ | 0.85 | 0.85 |

GNWT predicts that conscious perception should be marked by an ‘ignition’ event shortly after stimulus onset, as information from high-level visual areas (i.e., CS) updates the contents of the global workspace in PFC through long-range recurrent projections. After this initial ignition event, activity in PFC returns close to baseline. At stimulus offset, when the contents of consciousness change again, GNWT predicts that there should be another ‘ignition’ event in PFC—updating the contents of the workspace. The competitive dynamics at stimulus onset and offset are most naturally implemented in a dynamical systems framework by modeling PFC-to-PFC interactions as a winner-take-all system, which—following the initial competition between PFC masses—settles into a stable fixed-point attractor jointly constituted by the winning PFC mass and the CS mass. The GNWT-based model, therefore, consists of three strongly adapting neural masses: one representing CS and two representing the competitive interactions in PFC responsible for the update dynamics at stimulus onset and offset. The stimulus is delivered to CS, which then projects to the two PFC masses, which in turn compete in a winner-take-all mode to inhibit one another. After stimulus onset, strong adaptation reduces the activity of the winning (i.e., dominant) PFC mass. At stimulus offset, the previously suppressed mass (which is not subject to activity-dependent adaptation) then supersedes and inhibits the dominant mass, generating the hypothesized ignition update dynamics in PFC (cf. Fig. 1D[i]). To generate a single PFC time series, analogous to what would be observed in the MEG data, we averaged the simulated time series of the two PFC masses.

In contrast, according to IIT, conscious contents are maintained in posterior cortical structures throughout the duration of a conscious event via short-range reciprocal projections between low-level visual cortices (i.e., VIS) and higher-level category-selective (CS) cortices, generating synchronous activity between sensory regions, maintained for the duration of the stimulus presentation period (i.e., the duration of the visual experience). Within a dynamical systems framework, this prediction can be naturally read as implying that conscious contents are maintained in a high-activity stable fixed-point attractor jointly constituted by the VIS and CS masses. The IIT-based model, therefore, consists of two weakly adapting neural masses representing VIS and CS, respectively. The stimulus is delivered to VIS and is maintained in a high-activity attractor state jointly constituted by VIS and CS throughout stimulus presentation, as depicted schematically in Fig. 1D(ii).

#### 2.5.2 Quantitative model evaluation

We note that these models should not be regarded as direct neuronal implementations of IIT or GNWT; rather, they are simple phenomenological models (i.e., quantitative summaries) of the neural dynamics hypothesized to underlie conscious visual perception by each theory. These models allow us to quantitatively examine predictions about inter-areal coupling by applying FC measures identified in our data-driven analysis (with empirical MEG data) to our simulated neural data. Specifically, we compared the values of a given FC measure applied to model-simulated time series to those computed from empirically observed MEG signals using two measures: (1) the Kullback–Leibler (K-L) divergence [44] and (2) the Wasserstein distance [45]. For each model, we swept over the range of inter-areal coupling parameter values derived from stability analysis of each model (see Supplementary Methods, Sec. S1) and found the value of *W*_GNWT_ and *W*_IIT_ that best minimized the K-L divergence, giving each theory-based model the best chance we could to explain the observed data.

The K-L divergence, denoted *D*_KL_(*P* || *Q*), captures the amount of additional information (in units of bits) needed to reconstruct the probability distribution *P* when it is approximated by another distribution *Q*. This divergence is formalized as

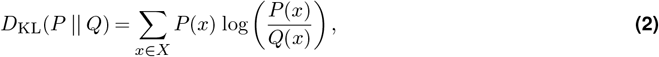

where *X* is the set of possible outcomes, and *P* (*x*) and *Q*(*x*) represent the probabilities assigned to each outcome *x* by distributions *P* and *Q*, respectively. Here, we use the K-L divergence to compute the information ‘cost’ of approximating the empirical distribution of FC values (*P*) with simulated distributions derived from each model (*Q*). In practice, we have *discrete* observations (i.e., the timepoints of the maximum squared Euclidean barycenter for empirical and model-simulated data) which we treat as independent, identically distributed (i.i.d.) samples *X*_1_, …, *X*_*n*_ and *Y*_1_, …, *Y*_*m*_ drawn from *continuous* distributions *P* and *Q*, respectively, with associated probability density functions *p* and *q*. Since the K–L divergence between continuous distributions is defined in terms of probability densities, here we estimate the KL divergence using the k-Nearest-Neighbor (kNN) estimator proposed by Wang et al. [46], which uses local neighborhood distance to infer relative probability densities of *p* and *q*. This estimator is given as

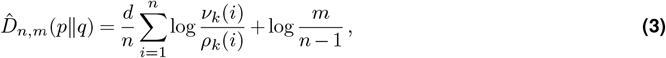

where *d* is the number of dimensions (in this case, *d* = 1), *ρ*_*k*_(*i*) is the Euclidean distance between *X*_*i*_ and its *k*-NN in {*X*_*j*_}_*j*_ ≠_*i*_, and *ν*_*k*_(*i*) is the distance from *X*_*i*_ to its *k*-NN in *Y*_*j*_. We applied this K-L estimator using an open-source Python implementation (https://github.com/nhartland/KL-divergence-estimators) that uses the `NearestNeighbors` function from *scikit-learn* [33]. Since the estimator from Wang et al. [46] is only asymptotically consistent, estimates are not guaranteed to be non-negative at a finite *n* (a setting in which local density estimates fluctuate), particularly when the *P* and *Q* distributions are highly similar. Negative estimates arise from finite-sample variability of the *k*-NN density estimator, even though the true divergence *D*_KL_(*P*∥*Q*) ≥ 0, so we treat negative values as zero in our interpretation.

The only parameter this estimator requires is the number of neighboring points to consider (*k*), which is important for the bias–variance tradeoff. As discussed in Wang et al. [46], a larger *k* generally yields lower variance but higher bias, though the bias can be mitigated through a larger sample size. By contrast, smaller values of *k* can yield unstable neighborhood geometry and divergent estimates at finite sample sizes, especially when comparing empirical data to dense simulated distributions. We examined a range of *k* values from [5, 35], finding that *k* = 20 was the smallest value for which both the IIT and GNWT models produced finite estimates across all evaluated parameters and stimulus presentation periods (with noise level *σ* = 1.0) for both raw and absolute value time series. The full range of *D*_KL_ values obtained in this sweep is shown in Fig. S1A. This choice is consistent with theoretical recommendations that 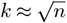 [47, 48]—*n* for each empirical *X* is 376, and 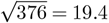 —and is further supported by the stability of parameter rankings across nearby values of *k*. After selecting *k* = 20, we also examined which inter-areal coupling strength magnitude (corresponding to *W* in Eq. (1)) minimized *D*_KL_ for each model. As shown in Fig. S1B, we computed the average *D*_KL_ between stimulus onset and offset for each *W* magnitude per model (i.e., *W*_IIT_ and *W*_GNWT_). The lowest average *D*_KL_ corresponds to *W*_IIT_ = 0.7 (*D*_KL_ = 0.13) and *W*_GNWT_ = 0.6 (*D*_KL_ = 0.08). Main results are therefore presented for *W*_IIT_ = 0.7 and *W*_GNWT_ = 0.6 with *k* = 20.

To complement the K-L divergence (which is not technically a distance metric and may be biased by outliers), we additionally computed the Wasserstein distance [45]—also known as the ‘earth mover’s distance’—which, in the discrete case, minimizes both the number of samples which must be moved and the distance those samples are moved for probability distributions *P* and *Q* to match (in units of total work). The K-L divergence with the *k*-NN estimator is sensitive to local density ratios, though the Wasserstein distance is not; as a geometric measure, it is more sensitive to how far the distribution mass in *Q* must be displaced to match the distribution of *P*. The Wasserstein distance (WD) was computed using the `wasserstein_distance` function from the *scipy* package (version 1.13.1) [32]. Together, the K-L divergence and the Wasserstein distance capture complementary aspects of alignment in density and geometry of the empirical and model-simulated results, respectively.

### 2.6 Code and data availability

All MEG data analyzed in this study are openly available upon registration at https://doi.org/10.17617/1.wqa3-wk71 [22]. All code needed to reproduce our analyses and visuals is freely available in our GitHub repository (https://github.com/anniegbryant/MEG_functional_connectivity) [49]. Preprocessed empirical data is also shared as a Zenodo repository at https://doi.org/10.5281/zenodo.18294156 [49].

## 3 Results

As depicted in Fig. 1A and described in the Methods (Section 2.2.1), we selected four key regions per hemisphere to represent areas hypothesized to underlie conscious visual perception by IIT and/or GNWT. These regions (in the 100-parcel resolution from Schaefer et al. [24]) are also listed in Table 1, and include the CS, VIS, PFC, and PAR areas of the cortex. We then comprehensively quantified MEG-derived FC between CS and each of the three other regions (taking the average time series across all epochs per event type and participant, also averaging the two hemispheres) using 246 bivariate time-series measures that span an interdisciplinary literature—from spectral properties to information-theoretic measures, using the `pyspi` (‘Python toolkit of statistics for pairwise interactions’) library [14]. Finally, we used the top-performing FC measures to analyze simulated time series from two theory-derived neural models, allowing us to interpret the empirical results in light of the pre-registered predictions of IIT and GNWT theorists as described in COGITATE Consortium et al. [11]. As described in the Methods (Sec. 2.4), our primary classification aim focused on stimulus-type decoding, and we explored two supplementary aims: domain-independent task relevance (‘Relevant’ vs. ‘Irrelevant’) and generalizability from one relevance type to another with the same pair of two stimuli. In all cases, we employed logistic regression classifiers with a grouped cross-validation strategy, reporting the mean *±* standard deviation (SD) out-of-sample accuracy (see Methods, Sec 2.4 for more details).

### 3.1 Multiple properties of inter-areal coupling can distinguish between stimulus types across region–region pairs

For the primary stimulus-decoding analysis, we first examined the overall range of cross-validated classification accuracy values across all 246 FC measures and experimental paradigm combinations (e.g., stimulus onset vs. offset, relevant non-target vs. irrelevant context). As shown in Fig. 2, many FC measures classified stimuli with 60-67% cross-validated accuracy across region–region pairs and task relevance settings (a more detailed breakdown by relevance type and stimulus presentation setting is shown in Fig. S2). This range sits within the context of prior visual stimulus-decoding analyses based on inter-areal connectivity derived from electroencephalography (EEG) time series [50] or combined EEG and functional magnetic resonance imaging (fMRI) data [51]. The presence of relatively high-performing FC measures in all stimulus pairs suggests there may be conserved signatures of neural activity in the functional pathways of interest while processing different visual stimuli. However, we note the marked variance in classification performance exhibited among FC measures across both region–region pairs and stimulus pairs (as shown in Fig. 2)—underscoring the heterogeneity in FC patterns across individuals, region–region pairs, and stimulus types.

**Figure 2.**
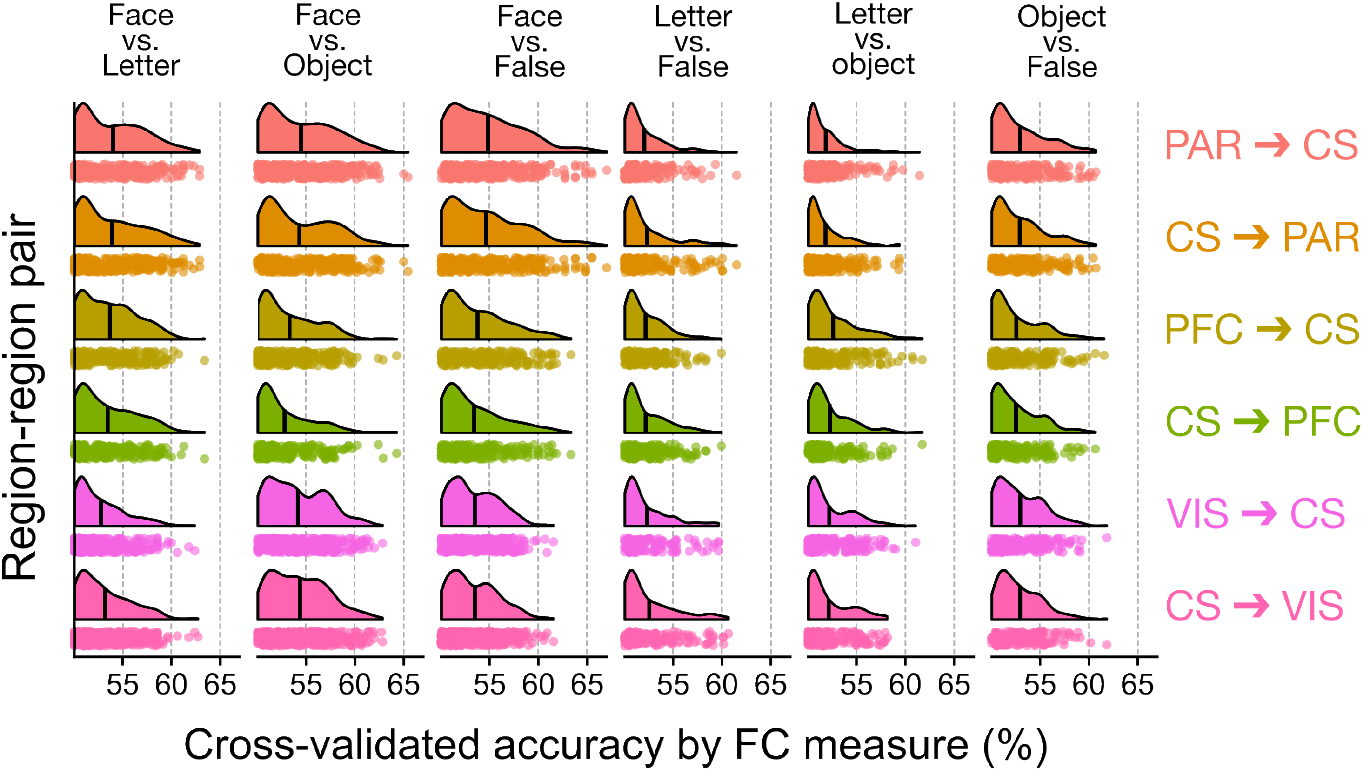
Many FC measures distinguish stimuli with ≥60% accuracy across region–region pairs, particularly for face stimuli. Each raincloud plot shows the distribution of classification accuracy values for each SPI in the corresponding stimulus pair (column) and region–region pair (row), partitioned by stimulus presentation period (‘on’ vs. ‘off’). Each dot represents one FC measure, and the vertical lines represent the mean cross-validated accuracy across all FC measures in the corresponding distribution. The *x*-axis was truncated at 50% accuracy, such that FC measures yielding lower than 50% accuracy are not shown here.

In the first supplementary classification aim, we probed the neural signatures of task relevance in a domain-general manner, testing whether any FC measure could distinguish ‘Relevant non-target’ versus ‘Irrelevant’ settings that combined all stimulus types. Surprisingly, none of the FC measures surpassed 56% accuracy with any region–region pair (cf. Fig. S3A), indicating that inter-areal coupling is less distinctive between domain-general task relevance settings. Our second supplementary classification aim was to test ‘cross-task’ transfer learning (as in COGITATE Consortium et al. [11]), in which we trained each classifier on FC measure values for a given stimulus pair (e.g., ‘face’ vs. ‘object’) during the ‘Relevant non-target’ setting (i.e., training on all N=94 participants) and tested the classifier in the ‘Irrelevant’ setting—and vice versa. In line with the predictions of GNWT, cross-task classification yielded similar accuracy distributions across FC measures and region–region pairs (including CS ↔ PFC) as in our primary stimulus type classification analysis (cf. Fig. S3B-C); therefore, we focus on the primary stimulus decoding classification analysis hereafter.

### 3.2 Statistics derived from the barycenter best distinguish stimuli, particularly faces

Having confirmed that multiple FC measures between the examined regions can distinguish between stimulus categories, we next turned our focus toward the highest-performing FC measures. This allowed us to make general inferences about the types of pairwise dynamics that contain the most information about consciously perceived visual stimulus categories. First, as shown in Fig. 3A, we identified the maximum classification accuracy across all FC measures, stimulus presentation periods (‘on’ or ‘off’), and category relevance settings (‘Relevant non-target’ or ‘Irrelevant’) for each stimulus pair (e.g., ‘Face’ vs. ‘Object’) and region–region pair (e.g., CS ↔ PFC). While we collapsed both directions into one region–region pair comparison each here for simplicity, the full results are shown in Fig. S4A. Across region–region pairs, the maximum classification accuracy values ranged between 62-67%, with all of the highest accuracy values corresponding to face stimuli and another category. However, we observed marked variability in classification performance across cross-validation folds (indicated by the shaded ribbons in Fig. 3A); therefore, while we report the mean accuracy *±* 1 SD, we draw inferences mainly from relative performance trends rather than absolute accuracy values.

**Figure 3.**
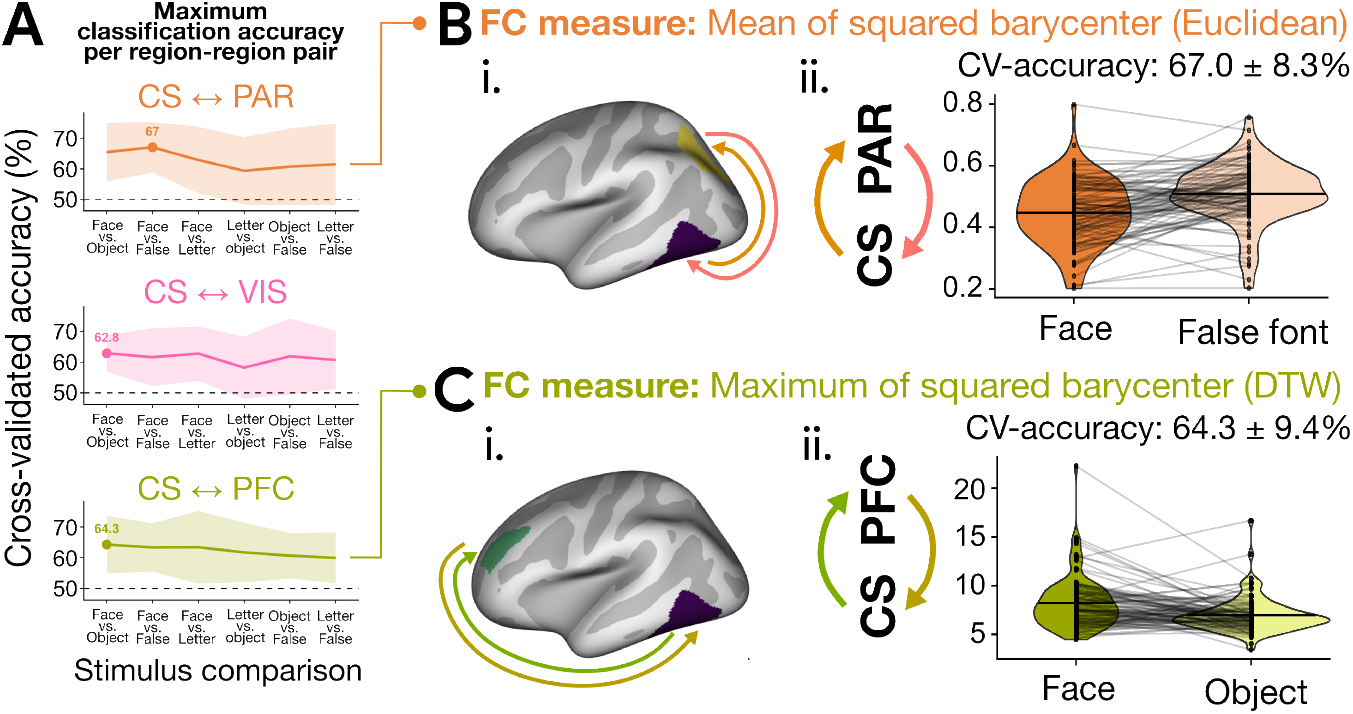
Highlighting the relatively strong performance of barycenter-based statistics across region–region pairs. **A**. For each connection between CS and another region, we plot the highest average accuracy with a line across all stimulus combinations (*x*-axis). Specifically, for each stimulus combination, the value indicates the peak cross-validated accuracy (with shaded ribbons indicating *±* 1 SD across test folds), corresponding to the top FC measure out of all 246 candidates—across either relevance condition (‘Relevant non-target’ or ‘Irrelevant’) and stimulus presentation period (‘on’ or ‘off’). For each region–region pair, the maximum overall accuracy is annotated. **B**. For CS ↔ PFC (i), the top-performing FC measure is the maximum of the squared barycenter with DTW averaging. (ii) This peak performance corresponds to ‘face’ versus ‘object’ during the task-irrelevant stimulus ‘on’ period (1000 ms). The violin plots show the distribution of this DTW-barycenter FC measure during ‘face’ versus ‘object’ trials across all N=94 participants. **C**. For CS ↔ PAR (i), the top-performing FC measure is the mean of the squared barycenter with Euclidean geometry. (ii) This peak performance corresponds to ‘face’ versus ‘false font’ during the task-irrelevant stimulus ‘on’ period (1000 ms). The violin plots show the distribution of this Euclidean-barycenter FC measure during ‘face’ versus ‘false font’ trials across all N=94 participants.

To better understand the types of dynamics captured by the top-performing FC measure classes, we focused on the top two classification measures overall, corresponding to ‘Face’ vs. ‘Object’ in the CS ↔ PFC regions (Fig. 3B) and ‘Face’ vs. ‘False font’ in the CS ↔ PAR (Fig. 3C), respectively. Of note, coupling between the two region–region pairs maximally distinguished stimulus categories based on the barycenter—a property originally developed in astrophysics to measure the center of mass between two orbiting celestial bodies [20], which extends here to capture the center of the pair of time series using a given distance minimization metric [52]. Also known as a Fréchet mean, the barycenter (*B*) of the activity in two regions is a new time series that minimizes the squared distances between the trial-averaged activity of those regions. For a pair of time series *x* and *y*, the barycenter *B* is computed as

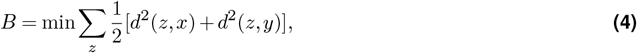

where *d* is the distance metric used to find the candidate barycenter (*z*) which minimizes the total squared distance between *x* and *y*. All barycenter-based statistics in `pyspi` [14] are implemented using the `tslearn` package [53], with three different types of geometry used to define the squared distances: Euclidean (in which the barycenter simplifies to the arithmetic mean of the two time series); dynamic time warping (DTW) averaging [52, 54], estimated with either the expectation maximization by default (prefix `bary_dtw` in `pyspi`) or with subgradient descent (prefix `bary_sgddtw` in `pyspi`) algorithms; and a differentiable variant of DTW geometry known as soft-DTW [55], which uses probabilistic phase alignment. We direct the interested reader to more detailed discussions about barycenter averaging with different distance metrics in Kathirgamanathan et al. [56]. Within each distance minimization metric (i.e., geometry type), we examined four output statistics from the `pyspi` library: the mean and maximum of the 1000 ms barycenter (corresponding to either stimulus onset or offset) and of the squared barycenter, respectively. The mean of the barycenter time series is sensitive to overall shared activity magnitude between the two regions sustained across the analysis window, while the maximum captures the strongest shared covariance from the baseline, offering sensitivity to transient, peak-like events (which can be obscured by averaging). Squaring the barycenter emphasizes larger-magnitude deviations from the center-of-mass signal of the two regions, which increases the sensitivity to transient high-amplitude co-fluctuations while reducing the influence of smaller co-variations. Summary statistics derived from the squared barycenter are more attuned to the presence and magnitude of joint activity between the two regions, regardless of activity sign.

In Fig. 3B-C, we compare the FC value distributions among all N=94 participants for the top-performing FC measure per region–region pair, such that the lines connect points corresponding to the same participant. For the CS ↔ PAR connection (Fig. 3B[i]), values are plotted for the mean of the squared Euclidean barycenter in Fig. 3B(ii) during ‘face’ versus ‘false font’ stimuli in the task-irrelevant setting. With the highest cross-validated accuracy across all evaluated comparisons (67.0 *±* 8.3%), this stimulus decoding was likely driven by the higher overall squared barycenter values throughout the entire 1000 ms presentation period for ‘false font’ versus ‘face’ stimuli. We schematically depict the computation of this FC measure using empirical CS and PAR time series from an example participant in Fig. S4B(ii). The time average of the squared Euclidean barycenter was nearly twice as high during ‘False font’ stimulus processing (0.59) than during ‘Face’ stimulus processing (0.30) for this example participant, which reflects greater sustained MEG fluctuation synchrony between the CS and PAR regions throughout the 1000 ms ‘False font’ stimulus period than during the ‘Face’ period.

For the CS ↔ PFC connection (Fig. 3C[i]), values are plotted for the maximum of the DTW barycenter in Fig. 3C(ii) during ‘face’ versus ‘object’ stimuli in the task-irrelevant setting. Unlike in Euclidean space (as in Fig. 3B), DTW geometry permits dilations and/or shifts across time for distance minimization between the two regions’ MEG activity. The maximum of the squared DTW-based barycenter provides insight into the variability in the underlying local activity in CS and PFC, as periods of higher MEG amplitude will be more heavily weighted in the temporal alignment and therefore exert greater influence on the barycenter. Indeed, the relatively high cross-validated accuracy of 64.3 *±* 9.4% is likely attributable to the generally higher maximum of the squared DTW barycenter during ‘face’ than ‘object’ stimuli, which suggests that the CS and PFC regions exhibited larger synchronized deviations from baseline activity during ‘face’ stimuli. The computation for this FC measure is schematically depicted using empirical CS and PFC time series from an example participant in Fig. S4C. The squared DTW-based barycenter peaked at a magnitude of 11.2 during ‘face’ stimuli—attributable to a sharp downward spike in both CS and PFC MEG amplitude at approximately 150 ms that is archetypic for MEG processing [57, 58]—compared to 7.5 during ‘object’ stimuli, in which MEG amplitudes were less variable and fluctuated on a slower timescale between CS and PFC.

### 3.3 Barycenter statistics dominate the top stimulus decoding results, along with basic covariance and spectral properties

In order to better understand the regional specificity of the top-performing FC measures, we next investigated the overlap in the number and type of FC measures that distinguished stimulus types well between region–region pairs. Since connectivity with the CS region is shared between IIT and GNWT, we focus on FC measures directed to and from the CS region, respectively—noting that undirected FC measures (such as the Pearson correlation coefficient) will yield bidirectionally symmetric values. We filtered the same classification results presented in Fig. 2 down to those with ≥ 60% mean cross-validated accuracy and tabulated the number of resulting FC measures projecting either from or to CS (e.g., from VIS, PFC, and PAR to CS). As shown in Fig. 4A-B, some FC measures uniquely distinguished stimuli in only one functional pathway; for example, 13 FC measures were unique to PFC → CS connectivity compared to that of VIS → CS or PAR → CS. However, we found three FC measures that distinguished at least one stimulus-pair to *and* from CS to all three other regions, all corresponding to the maximum of the squared barycenter computed with different geometries for the distance metric (Euclidean, DTW, and a variation on DTW geometry known as soft-DTW [55]). This suggests that the maximum of the squared barycenter captures stimulus-relevant information (for specific stimulus types) in all functional pathways relevant to IIT and/or GNWT, poising it as a useful type of measure with which the theories may be quantitatively evaluated and contrasted.

**Figure 4.**
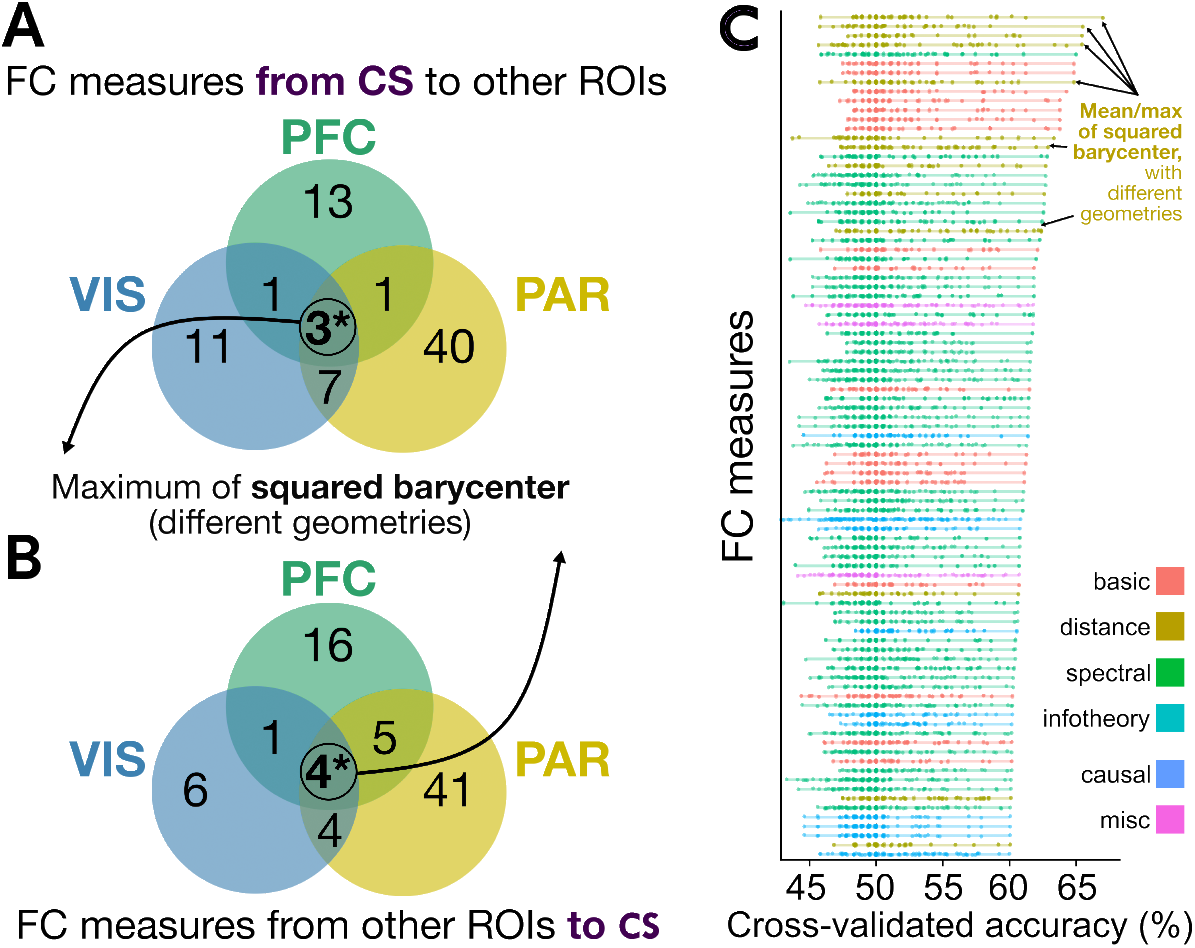
The mean and maximum of the squared barycenter, computed using distance metrics, generalize as top-performing FC measures for stimulus decoding across region–region pairs. **A**,**B**. Here, the number of FC measures yielding at least 62% cross-validated classification accuracy in one or more stimulus pairs is tabulated for each region as either the source (**A**) or target (**B**) region for the corresponding interaction. Ellipses within the Venn diagram are color-coded to indicate the corresponding region, following the brain maps depicted in Fig. 1A. The circled ‘3’ at the center of the Venn Diagram in **A** indicates that three FC measures distinguished between one or more stimulus pairs with ≥60% accuracy from CS to all other regions. These three FC measures are also contained in the ‘4’ at the center of the Venn Diagram in **B**, comprising the FC measures with ≥60% accuracy from all other regions to CS. The three measures shared between **A** and **B** all correspond to the maximum of the squared barycenter derived using different geometries. **C**. For each of the 95 FC measures summarized in **A-B**, the cross-validated classification accuracy values are plotted for all 144 parameter combinations (region–region pairs, stimulus comparisons, onset vs. offset, and relevance settings). FC measures are ranked in descending order of the maximum accuracy and colored by literature category, with the legend indicated at the bottom right.

All FC measures can be categorized into one of six literature-based domains (‘basic’, ‘distance’, ‘spectral’, ‘info-theory’, ‘causal’, or ‘miscellaneous’; cf. Cliff et al. [14]), so we next compared the FC measure type composition for the top-performing measures shown in Fig. 4A-B. As shown in Fig. S5A-B, undirected FC measures corresponding to distance measures (including the barycenter) and frequency-based coupling (e.g., power envelop correlation) comprised the top-performing statistics for all three functional pathways both to and from the CS region. Although there is high-level similarity among FC measures that distinguish stimuli across these functional pathways, the relatively low overlap shown in Fig. 4A-B indicates that the particular FC measures differ between region–region connections. The overlap between CS ↔ VIS and CS ↔ PFC corresponds to the mean barycenter computed with soft-DTW geometry, while the FC measure overlap between CS ↔ PFC and CS ↔ PAR is driven predominantly by spectral properties like power envelop correlation and imaginary coherence (more detail on these FC measures is shown in Fig. S5C-D). In addition, statistics capturing basic covariance structure (including the Pearson correlation coefficient) comprised many of the top-performing FC measures between CS ↔ PFC and CS ↔ PAR, but not CS ↔ VIS.

Beyond characterizing the maximum decoding accuracy for each FC measure across region–region pairs (which represents a ‘best-case’ two-class stimulus discrimination scenario), we also examined the full performance range for each of the 95 FC measures contained in Fig. 4A-B (i.e., all FC measures with at least ≥ 60% cross-validated accuracy). Fig. 4C shows all decoding accuracy values per FC measure, tabulated across 144 combinations of stimulus contrasts, region–region pairs, task relevance settings, and presentation periods (onset vs. offset). The highest maximum accuracy values correspond to the mean and maximum of the squared barycenter derived from different geometries, as annotated in Fig. 4C—though even these FC measures fail to surpass chance classification performance (50%) in many settings. This visualization demonstrates the variability in decoding accuracy exhibited by all 95 examined FC measures, none of which outperformed chance in all 144 experimental parameter combinations.

### 3.4 Testing predictions via data-driven analysis of theory-driven neural modeling

In the previous sections, we identified a multitude of FC measures that met our core stimulus decoding classification aim across region–region pairs. In particular, FC measures derived from the barycenter generalized across all of the relevant region–region pairs relevant to both IIT and GNWT. Barycenter-based measures are fast to compute, simple to interpret, and novel to functional neuroimaging—prompting further investigation with modeling. To derive theoretical insight into the neural processes predicted to underlie conscious vision using barycenter-based FC, we constructed two neural mass models [21] tailored to recapitulate each theory’s pre-registered predictions [11] for neural activity in each region (for details see Methods Sec. 2.5 and the schematic depiction in Fig. 1D).

Importantly, because each theory generates predictions about coupling between different region–region pairs (CS ↔ VIS for IIT and CS ↔ PFC for GNWT), our goal in this section is to evaluate the relative success (rather than the overall variance explained) of each theory-derived model in approximating the empirical data. The models are stimulus-agnostic and are built to reproduce the empirical Euclidean barycenter values averaged over all stimulus types. We focused on the stimulus-averaged dynamics rather than the stimulus-specific dynamics, as theorists from neither GNWT nor IIT provided stimulus-specific predictions. Rather, they specified predictions about the qualitative features of inter-areal coupling (e.g., synchronized activity between CS and VIS) hypothesized to underlie conscious perception. In addition, to avoid a mismatch between simulated and empirical barycenter values caused by changes in dipole polarity across regions (cf. Fig. S6A-B), we specifically computed the absolute value (i.e., magnitude) of both the empirical and simulated time series before computing the Euclidean barycenter in this section. As a robustness test, we show in Fig. S7 that the stimulus classification results based upon the squared barycenter (across all four possible distance metrics) are not systematically biased by this extra preprocessing step.

Our model simulation process is schematically depicted in Fig. 5A. Using the IIT- and GNWT-based models, we simulated 1000 trials with dynamical noise and then subtracted the average neural activity before stimulus onset (i.e., the ‘baseline’) from each trial (Fig. 5A[i-ii]). We approximated noise in the measurement process by adding zero-mean independent Gaussian noise to each trial (Fig. 5A[iii]). Each trial was divided into 1000 ms segments corresponding to stimulus-on and stimulus-off periods for analysis. We show results obtained using *σ* = 1.0 added noise in the main text, with results from the full range of 0.5 ≤ *σ* ≤ 1.0 depicted in Fig. S8. Within each stimulus-on and stimulus-off epoch, we computed the maximum of the squared barycenter across Euclidean and non-Euclidean geometries (Fig. 5A[iv]). For clarity, we show the results for the maximum of the squared barycenter derived from Euclidean geometry; results are qualitatively comparable using DTW, soft-DTW, and subgradient descent (SGD-)DTW geometries for the distance metric (cf. Fig. S9).

**Figure 5.**
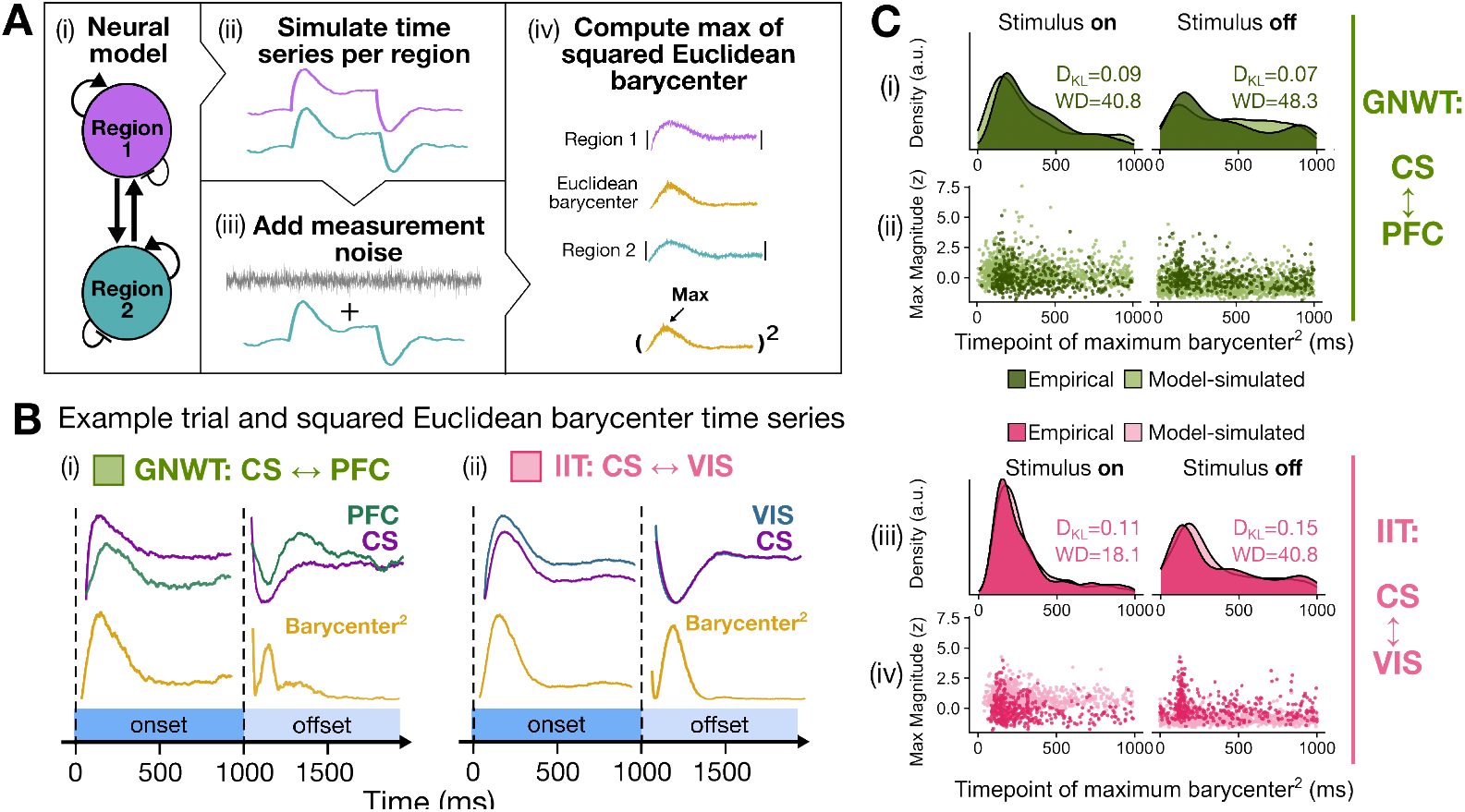
Comparing model-based predictions for barycenter with results from empirical COGITATE MEG data. **A** Graphical illustration of the model analysis pipeline. For both models, we simulated 1000-trials with noise, and after baseline subtraction and added zero-mean Gaussian noise to each time point of the simulation to mimic the effect of measurement noise before computing the max of the squared barycenter for each trial. For comparability with the empirical data, the barycenter was computed after taking the absolute value of the time series, thereby avoiding issues of dipole polarity reversal across regions. **B**. Illustration of single-trial neural activity and barycenter time series for GNWT- and IIT-based neural mass models. Because of the offset between the CS and PFC masses, and the winner-take-all dynamics, the timing of the peak barycenter values is more variable across trials in the GNWT-based model than in the IIT-based model. **C**. Here, we examine the timing and magnitude of the squared Euclidean barycenter maximum in both empirical and model-simulated time series (with noise *σ* = 1) for both IIT and GNWT during stimulus onset and offset. First, the probability distribution for the timing of the barycenter maximum for each (i) CS↔PFC and (iii) CS↔VIS bivariate time series is summarized via *k*-NN density estimation for each stimulus presentation period and theory. The empirical density distribution (kernel-estimated only for visualization purposes) is dark-shaded, and the model-simulated density distribution is light-shaded in each case. The Kullback-Leibler divergence [44], *D*_*KL*_`(Empirical` || `Model)`, is shown for each theory and stimulus presentation period, in units of bits, along with the Wasserstein distance (WD) in units of total work. (ii, iv) The same maximum squared Euclidean barycenter values comprising the density plots in (i) and (iii) are plotted, with the maximum timing on the *x*-axis and the magnitude of the corresponding squared Euclidean barycenter on the *y*-axis. The same color scheme is used as in (i) and (iii).

Briefly, as schematically represented in Fig. 5B(i), GNWT predicts that visual consciousness is dependent on neural signal propagation from CS to PFC regions, leading to a late ‘ignition’ of the global workspace after stimulus onset and again at stimulus offset (we show in Fig. S6F(ii) that the PFC does indeed show a secondary bump in activity after stimulus onset, in line with the predictions of GNWT theorists). The GNWT-based model has two key features responsible for the ignition-like updating dynamics in PFC at stimulus onset and offset: (1) the winner-take-all competition between the two PFC masses, and (2) the slow timescale of the PFC neural masses (*τ*_PFC_=80 ms) relative to the comparably fast CS neural mass (*τ*_CS_=40 ms). This leads to a peak in PFC activity ∼ 200 ms after stimulus onset—approximately 150 ms after the peak in CS activity, which peaks ∼ 50 ms after stimulus onset.

In terms of the (squared) barycenter, which tracks the center of mass of the two time series, the offset in peak activity between the two masses reduces the amplitude of the (squared) barycenter. The relatively low amplitude of the peak PFC signal also means that in the presence of measurement noise, the maximum of the barycenter is broadly distributed throughout the stimulus-on epoch. At stimulus offset, both the CS and PFC masses show a brief dip in activity due to the accumulated adaptation current. The winner-take-all competition between the two PFC masses then generates a secondary bump of activity approximately 300 ms after stimulus offset, at which point the previously suppressed mass takes over. Similar to the stimulus-on period, the offset in the peak activity between the CS and PFC masses, combined with measurement noise, yields a broad distribution of maximum (squared) barycenter values across the stimulus-off period.

In contrast, as schematically represented in Fig. 5B(ii), IIT predicts that the conscious experience of a visual stimulus involves synchronized and sustained activity between CS and VIS (i.e., the ‘posterior hot zone’) throughout stimulus presentation. The key features driving the dynamics of the IIT-based model are the reciprocal connectivity between the VIS and CS masses and the comparably fast and similar timescale of the two masses (*τ*_VIS_ = 20 ms, *τ*_CS_ = 40 ms). Because of the similar timescale and reciprocal connectivity, both masses increase (and decrease) synchronously following stimulus onset and offset. During the stimulus presentation period, activity is maintained in a stable high-activity attractor state. The synchronized peaks and troughs in activity at stimulus onset and offset generate large peaks in the (squared) barycenter amplitude, yielding a high-density concentration of maximum timing values at the beginning of the epochs across trials.

Qualitative comparison of the empirical and simulated distributions shows that both the GNWT- and IIT-based models roughly approximate the empirical MEG data, a result reflected in the quantitative comparison of the simulated and empirical distributions of maximum squared Euclidean barycenter timings with the GNWT-based model and IIT-based model providing comparable approximations to the empirical data for both stimulus-onset (KL_GNWT_ = 0.09 nats vs. KL_IIT_ = 0.11 nats) and offset (KL_GNWT_ = 0.07 nats vs. KL_IIT_ = 0.15 nats) periods. This result is also reflected in the Wasserstein distance at stimulus-onset (WD_GNWT_ = 40.8 vs. WD_IIT_ = 18.1) and stimulus-offset (WD_GNWT_ = 48.3 vs. WD_IIT_ = 40.8). In other words, the maximum squared barycenter values are well approximated by both theory-derived models in terms of both information cost (K-L divergence values) and Wasserstein distance. Importantly, this pattern is robust across various noise levels (cf. Fig. S8) and task relevance settings (cf. Fig. S10). This conclusion notably depends on the model-based implementation of the theoretical predictions and may be subject to revision. In the context of this caveat, the predictions of both theories seem to accurately describe empirical patterns of connectivity, suggesting that while both theories can be cast as mutually exclusive, their predictions are not necessarily in disagreement.

## 4 Discussion

Here, we present an approach for data-driven identification, evaluation, and theoretical interpretation of connectivity-based neural correlates of conscious visual perception. Our work is built upon the datasets and preregistered hypotheses generated by the COGITATE Consortium adversarial collaboration that tested the predictions of IIT and GNWT [11]. We focused here on functional connectivity (FC), intending to find a measure(s) that provided reliable stimulus-type decoding—reasoning that if a neural process underlies the content of conscious visual perception, variation in that process should encode information relevant to these contents. Through the most comprehensive comparison performed to date, we systematically evaluated 246 different measures of pairwise coupling, derived from an interdisciplinary time series literature [14], computed between regions hypothesized to drive conscious vision in IIT and/or GNWT, and used theory-driven neural modeling to derive theoretical insight into the processes hypothesized to underlie conscious vision by each theory.

For each region–region pair, we identified multiple FC measures that decoded stimulus type with >60% cross-validated (out-of-sample) accuracy. While relatively modest in magnitude for binary classification, the higher-performing accuracy range of 60–67% is broadly consistent with prior stimulus-decoding work based on inter-areal FC derived from EEG or combined EEG–fMRI data [50, 51]. We note that direct comparison with MEG-based FC decoding studies is challenging, as most MEG decoding analyses focus on within-region signals at specific time points; by contrast, our approach summarizes inter-regional connectivity over extended stimulus periods (1000 ms) to prioritize generalization. The raw accuracy values should also be interpreted within the context of our intentionally conservative analysis framework. Specifically, we applied rigorous cross-validation and performed decoding across (rather than within) individuals, both of which are expected to yield lower (but more generalizable) performance estimates in a heterogeneous population [59]. Together with the strong correspondence between cross-validated accuracy and AUC (Pearson’s *R* = 0.71), the consistency across cross-validation folds and classification problems suggests the presence of a reliable and systematic stimulus-relevant signal.Rather than simply seeking to maximize decoding accuracy, our objective in this work was to leverage the rich interdisciplinary time-series literature implemented in the `pyspi` library [14], to identify the types of coupling properties that generalize across stimulus types and participants. The top-performing FC measures included those commonly used to quantify activity from MEG data, such as the power envelop correlation [17, 18]. However, our data-driven approach also uncovered previously unexplored FC measures that meaningfully distinguished stimulus conditions across region–region pairs, such as the barycenter derived from both Euclidean and non-Euclidean geometry. For example, the top-performing FC measures that yielded >60% stimulus decoding accuracy across region–region pairs common to both GNWT (CS ↔ PFC) and IIT (CS ↔ VIS) were based upon the maximum of the squared barycenter. The barycenter was originally developed in astrophysics to track the center of mass between two orbiting celestial bodies [20], and is sensitive to both the amplitude and relative phase of each time series (especially when computed using Euclidean distance). Interpreted through the lens of FC here, barycenter-derived measures capture properties of the central alignment of two regions’ dynamics using a given distance minimization metric [52]. In particular, the maximum of the squared barycenter identifies the strongest distance metric-defined covariance between the two regions over a given 1000 ms stimulus presentation window; both the magnitude and the timing of this value can be meaningfully analyzed. Since this class of features maximally decoded stimulus types, we reasoned that statistics derived from the barycenter form strong candidates for encoding information relevant to conscious visual perception. To our knowledge, this is the first application of barycenter-derived statistics to FC in the human brain, underscoring the utility of a data-driven framework to derive new informative measures for a given use case.

After identifying these barycenter-based FC measures, we next sought to derive theoretical insight into the neural processes hypothesized to underlie conscious vision by each theory by comparing empirical barycenter values to simulated barycenter values derived from two simple neural mass models [21] tailored to recapitulate the pre-registered predictions about regional activity by IIT and GNWT theorists (see COGITATE Consortium et al. [11]). For the GNWT-based model, the winner-take-all competition between the two PFC masses, coupled with their slow timescale, yielded a broad distribution of maximum (squared) barycenter timings across the stimulus onset period and stimulus offset period. In contrast, in the IIT-based model, the CS and VIS masses exhibit a similar timescale and strong reciprocal connectivity, such that both masses increased (and decreased) synchronously at stimulus onset and offset. This generated tightly peaked distributions of maximum squared barycenter values at the beginning of each onset and offset period. Quantitatively, simulated barycenter values from both the GNWT-based model and the IIT-based model provided comparable approximations to the empirical data in terms of both information cost (lower K-L divergence values [44]) and the Wasserstein distance (also known as ‘earth mover’s distance’ [45]).

While the two theories are cast as mutually exclusive by their advocates, when the predictions are translated into models that can be quantitatively compared to data, it can be argued that the predictions themselves are not mutually exclusive. Although the predictions of both theoretical camps provide comparable approximations to the empirical data, IIT theorists predicted that ignition-like dynamics in the PFC should reflect task demands rather than conscious perception. In contrast, GNWT theorists remained agnostic about the role of synchrony in the sensory cortices. The success of the GNWT-based model, which includes ignition-like WTA dynamics, is contrary to the predictions of IIT theorists, whereas the success of the IIT-based model is not necessarily in conflict with the predictions of GNWT theorists. Again, however, we note that this conclusion is conditional upon the model-based implementation of hypotheses from GNWT and IIT, and may be subject to revision should another model—still consistent with the hypotheses—be put forward.

The framework and findings in this work should be considered alongside technical and conceptual limitations that inspire future investigation. For example, even among the top-performing FC measures (including statistics derived from the barycenter), there is variability in stimulus decoding performance across different experimental parameters. Much like the ‘no free lunch’ theorem in machine learning [60, 61], our findings do not suggest a singular FC measure that will universally perform well across all stimulus decoding contexts. It remains to be seen, therefore, whether statistics based upon the barycenter generalize as top performers beyond the current context. There are also potential limitations specific to our MEG-based connectivity analyses. For example, we averaged across all epochs within a given event-related field type for each participant, such that each of the N=94 participants was represented by a single data point per FC measure in a given classification analysis. While the temporal averaging was performed to enhance the signal-to-noise ratio and reduce computational load [30], this design might also obscure relevant inter-trial differences and/or inter-individual heterogeneity in neural responses to the same stimulus type and experimental setting. Additionally, we employed the same preprocessing pipeline as in COGITATE Consortium et al. [11], which employed the widely-used MNE-Python workflow [62]. However, issues remain regarding proper source localization, environmental/biological artifact removal, and latency variability, the latter of which poses issues for interpreting asynchronous (e.g., lagged) FC measures [63]. Field spread is also a concern due to the difference in spatial proximity between regions—for example, VIS and CS regions are much closer together than VIS and PFC—which also means that VIS and CS regions are more likely to exhibit similar MEG time series due to spatial autocorrelation that decays with distance [64].

We also note that our models are phenomenological (rather than mechanistic) and should be thought of as quantitative summaries of the neural dynamics hypothesized to underlie conscious perception by IIT and GNWT—rather than full implementations of the theories themselves. In addition, we note that models should ideally be built before data is collected. Indeed, a key limitation of the current study is that we built the models post-hoc based upon preregistered *qualitative* hypotheses. Because these hypotheses are specified in words rather than equations, there will always be an imprecise mapping between formal models and the hypotheses themselves. It is therefore critical for the field to move towards formulating hypotheses in quantitative terms that can directly link to formal models—which provide a concrete starting point for revisions that may better explain the observed empirical data. This is especially important when predicted time-series features are not identified as informative. If a model includes the full suite of neural processes hypothesized by a theory to underlie a cognitive/perceptual process of interest, the model can still be used to inform the interpretation of other time series features that are identified as informative.

We fixed the parameters of each model by making a series of severe assumptions about the nature of the effective dynamics, which eliminated all free parameters except for the strength of inter-areal connectivity. We then used stability analysis to identify the range of parameter values consistent with the hypothesized dynamics, taking the parameter value that best minimized the difference between simulated and empirical data. An important next step in the analysis pipeline will therefore involve leveraging previous work on model fitting and model comparison to fit and evaluate families of theory-derived models, which include a broad range of theory-relevant parameters [65, 66]. Finally, we note that because the experimental paradigm targeted the mechanisms of conscious vision and did not contrast conscious and unconscious vision, the study should be viewed as strictly testing the predictions of IIT and GNWT theorists in the domain of conscious vision. The results, therefore, do not bear strongly on the question of whether the mechanisms posited by each theory are necessary or sufficient for perceptual consciousness.

In practice, theory testing (and refinement) is an iterative, rather than absolute, process [67, 68]. The predictions set out by the COGITATE Consortium [11] are not derived from the core of IIT or GNWT, and therefore may be revised whilst leaving the core tenets of each theory intact [67]. The predictions should be viewed instead as those of specific readings of each theory, rather than predictions of the theories themselves in some absolute sense. There are many steps and auxiliary assumptions that must be made to connect the core of a theory to data. This is not to say that the accuracy of the predictions should not increase or decrease our credence in each theory [68]; rather, we point out that we should not expect to falsify IIT or GNWT outright based on one experiment [69]. There is, therefore, room for productive disagreement between researchers even within the same theoretical camp [70, 71] about the best way to connect a theory to data. We should not expect perfect agreement about the best way to do this. Rather, we argue that the most productive way forward for each theory is to make the differences in assumptions explicit and quantitatively compare models derived from the same theory under the divergent assumptions as we have done here for the predictions set out by the COGITATE Consortium [11].

## 5 Conclusions

We present a roadmap for identifying and evaluating candidate neural correlates of conscious visual perception in a data-driven manner that integrates with theory-driven modeling to derive quantitative and interpretable insights. Importantly, the data-driven nature of this approach means that one need not make *a priori* assumptions about which time series feature(s) will be best suited for the process of interest. Once informative time-series features are identified, modeling plays an indispensable role in mapping between the neural processes hypothesized to underlie the observed dynamics and the empirically measured time-series features themselves. The models we specified here are deliberately as simple as possible, prioritizing interpretability, and are also specifically constructed to simulate Euclidean barycenter values, which are sensitive to the relative amplitude and phase of the two time series.

To our knowledge, this is the first study to systematically compare hundreds of FC measures for decoding visual stimuli across task relevance settings. We focused here on bivariate coupling measures using the `pyspi` toolbox [14], though this approach is readily extendable to highly comparative analysis of both univariate time series [72–74] (to focus on local dynamics of a single region) or higher-order interactions comprising three or more regions, a rapidly growing area of research [75, 76]. Combining across different scales of dynamics, from local to higher-order, could paint a fuller picture of the rich neural complexity driving a biological phenomenon like conscious visual perception [15]. In addition, an important avenue for future work will be to quantitatively compare measures that distinguish between conscious and unconscious stimuli, allowing us to isolate the processes that make a stimulus conscious—rather than focusing on the neural process underlying consciously perceived stimuli, as we did here. Importantly, our data-driven approach also generalizes across neuroimaging modalities and, when combined with neural modeling, offers clear insights into the processes hypothesized to generate empirically observed brain dynamics.

Large openly available datasets are increasingly common in consciousness science; concurrently, the field is progressively moving towards more quantitative neural theories [77–82]. There is, therefore, a unique opportunity for collaborations between experimentalists, data analysts, and neural modelers to systematically build, and quantitatively compare, measures and theories of consciousness using data-driven but theoretically interpretable frameworks such as the one we present here.

## 6 Acknowledgments

We are grateful to the COGITATE Consortium for their Herculean effort in setting up the adversarial collaboration and for making the data openly available. In addition, we would like to thank our supervisors, Ben Fulcher and Mac Shine, for their guidance and encouragement throughout this project. We would also like to thank Borjan (Boki) Milinković, Niccolo Negrò, and the members of the Mudrik lab for their detailed and thoughtful feedback on the work. We acknowledge the use of high-performance computing clusters at The University of Sydney (School of Physics cluster and University-wide Artemis system) in generating results for this paper. AGB acknowledges support from the Research Training Program (Australian Government) and the Paulette Isabel Jones Career Award (The University of Sydney). CJW acknowledges support from the DVCR Strategic Postgraduate Research Scholarship (The University of Sydney).

## S1 Supplementary methods

### S1.1 Model derivation

The models described in the main text represent the effective dynamics of the Wilson-Cowan neural mass model of excitatory (*r*_*E*_) and inhibitory (*r*_*I*_) cortical dynamics [38] under the slow inhibitory pressure of spike-frequency adaptation (*a*):

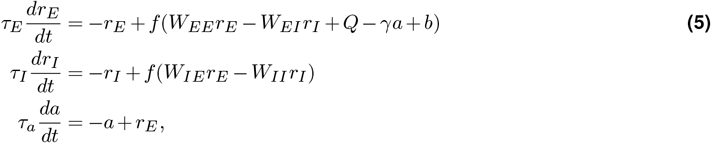

where *f* (*x*) is the nonlinear (typically sigmoidal) population transfer function; *W*_*ij*_ denotes the connection strength between pre- and post-synaptic cells (i.e., *i* ← *j*); *τ*_*E*_, *τ*_*I*_, and *τ*_*a*_ denote the excitatory and inhibitory neuronal time constants and adaptation time constant; *γ* denotes the strength of the hyperpolarising adaptation current; *Q* denotes the external current entering the population (i.e., input from an external stimulus or input from the surrounding network); and *b* denotes the baseline levels of excitation due to arousal.

Leveraging the physiological observation that GABAergic neurons respond rapidly ([41]; i.e., *τ*_*E*_ ≫ *τ*_*I*_), we approximated the inhibitory population with its equilibrium value *r*_*I*_ ≈ *r*_*I*_ (∞), which—assuming the inhibitory population is in the linear region of the transfer function—is given by

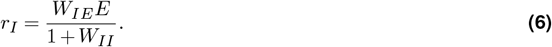

Under the assumption that the currents entering the excitatory population are balanced (such that *W*_*EE*_ − *W*_*EI*_ *W*_*IE*_*/*(1 + *W*_*II*_) = 0), substitution of Eq. (6) into Eq. (5) leads to the following simplified expression for the effective dynamics of *r*_*E*_ (where we have dropped subscripts for ease of notation):

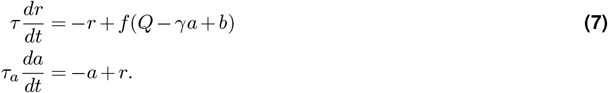

Under the assumption that both populations are operating in the linear region of their transfer functions, we replaced the typical sigmoidal transfer function with the threshold linear function *f* = max(*x*, 0).

### S1.2 Parameter selection

As described in the main text, we implemented each theory’s predictions in neural models by associating the qualitative predictions with attractor dynamics that we could then use to constrain the parameters of each neural model via standard stability analysis [37, 83]. To restrict the number of parameters, we assumed that all weights are symmetrical (*W*_*ij*_ = *W*_*ji*_). Values for *τ* were selected from the empirical literature on the timescale of each region [42]. This leaves the IIT-based model with three free parameters (*W*_IIT_,*γ,b*) and the GNWT-based model with four (*W*_GNWT_,*W*_P_,*γ,b*). Below, we derive fixed values for *W*_P_,*γ,b*, and the interval for *W*_THEORY_ over which each model is stable.

### S1.3 IIT-based model connectivity parameters

Based on the prediction of the COGITATE Consortium [11] that under IIT, conscious contents should be maintained in posterior cortical structures throughout the duration of a conscious event via reciprocal projections between visual (i.e., VIS) and category-selective (CS) cortices, we built a model consisting of two reciprocally connected weakly adapting neural masses representing VIS (*r*_*V*_) and CS (*r*_*C*_), respectively.

In order to ensure that both masses were active and stable for the full duration of the stimulus presentation period, in line with predictions for IIT in COGITATE Consortium et al. [11], we searched for the region of parameter space compatible with a stable fixed point. As we wanted the model to be at a stable fixed point for the full duration of the stimulus presentation period, we set adaptation to its equilibrium value *a* = *a*(∞) = *r*. This ensured that the population would remain at a stable fixed point even under the inhibitory pressure of adaptation, and reduced the four-dimensional system of equations to two:

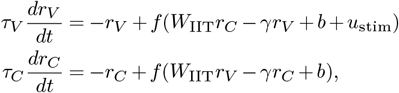

where *W*_IIT_*r*_*C*_ and *W*_IIT_*r*_*C*_ are the inter-areal connections between each population, and *u*_stim_ is the external stimulus injected into the VIS population (in Eq. (5), these terms were grouped together under the external current term, *Q*). During stimulus presentation (*r*_*V*_, *r*_*C*_ *>* 0), which—given the simple activation function *f* = max(*x*, 0)—reduces to

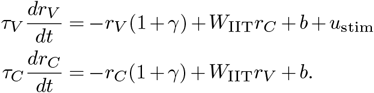

with Jacobian matrix:

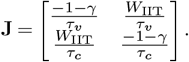

A stable fixed point requires that the trace of the Jacobian is negative (tr(**J**) *<* 0) and the determinant is positive (det(**J**) *>* 0). A negative trace requires that

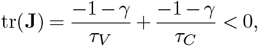

which is satisfied for all physiologically plausible values of *γ, τ*_*V*_, and *τ*_*C*_, which are always positive. The conditions for a positive determinant are given by,

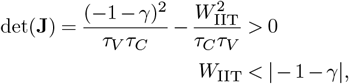

which provides us with a parameter interval for *W*_IIT_. As simulations were performed in the presence of noise, and *τ*_*a*_ ≫ *τ*_*V*_, *τ*_*C*_, simulations were run on the restricted interval 0.1 ≤ *W*_IIT_ ≤ 0.9 which satisfies the above inequality even when the contribution of *γ* is negligible (i.e., at the start of each stimulus presentation period), and is robust to noise induced fluctuations in firing rate.

### S1.4 GNWT-based model connectivity parameters

Based upon the predictions outlined by the COGITATE Consortium [11], along with previous relevant neuronal modeling [80] and theory [84], we implemented the GNWT-based model in a network of three strongly adapting neural masses: one representing CS and two representing within-prefrontal cortex (PFC) interactions. To generate ignition-like dynamics, we imposed winner-take-all (WTA) conditions on the PFC-to-PFC connections, leading to the suppression of one PFC mass at the expense of the other. To ensure the stability of the WTA regime over the full stimulus presentation period, we imposed conditions for a stable fixed point on the connections between PFC and CS.

We start by deriving the conditions necessary for WTA PFC dynamics. Both PFC populations (*r*_*P* 1_,*r*_*P* 2_) receive driving projections from the CS mass, which, for simplicity, we treat as a constant (again denoted by *Q*). Again, we assume that adaptation has reached its equilibrium value *a* = *a*(∞) = *r*, as we want to obtain WTA conditions valid for the full stimulus presentation period. This leaves us with the following two-dimensional system describing within-PFC dynamics:

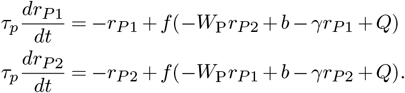

For *r*_*P* 1_, *r*_*P* 2_ > 0 the Jacobian matrix is given as

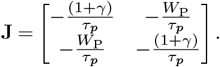

WTA dynamics require the fixed point for *r*_*P* 1_, *r*_*P* 2_ *>* 0 to be a saddle node, which occurs when the determinant of the Jacobian is negative,

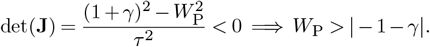

For robust WTA behavior in the face of noise, we want *W*_P_ to have a value that is well above the bounds of the inequality. In addition, once a population is suppressed, as long as it remains suppressed, the degree of suppression does not alter the dynamics. We therefore used a fixed value of *W*_P_ = 2.1 and found the range of CS-to-PFC connection strengths over which the WTA regime was stable.

If the model is in a WTA regime, one of the PFC masses is suppressed, reducing the GNWT-based model to a two-dimensional system describing the interaction between the dominant PFC mass and the CS mass:

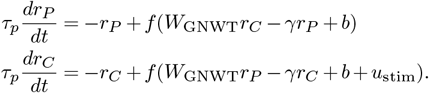

The conditions for a stable fixed point are the same as they were for the IIT-based model, resulting in *W*_GNWT_ *<* | − 1 − *γ*|. Because the WTA regime within the PFC population is sensitive to noise-driven fluctuations in firing rate, we restricted the model to the slightly narrower weights interval 0.3 ≤ *W*_GNWT_ ≤ 0.85.

### S1.5 Adaptation and arousal parameter selection

We settled on fixed values for *b*, which were shared across models, and *γ*, which differed between models, by calculating the equilibrium values of each system and ensuring that the resulting values qualitatively matched the predictions of each theory set out by the COGITATE Consortium [11].

For non-zero firing rates, both models can be written as a linear system of the form

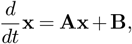

which has an equilibrium at **x**_eq_ = **A**^−1^**B**. Here, **A** and **B** are given by

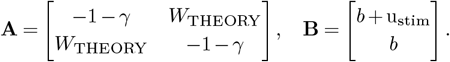

For simplicity, we set u_stim_ = 1, and *b* = 0.85, such that each model had a non-zero baseline firing rate given by **x**_eq_ across the full range of *W*_THEORY_ values, well above the rectification threshold of *f* to justify the assumption of linearity.

We then chose the values of *γ* for the GNWT-based model to be as large as possible, resulting in a strong reduction in firing rate post-stimulus onset (in line with predictions from the COGITATE Consortium [11]) whilst still satisfying the inequality for WTA activity, resulting in *γ* = 0.85. We similarly chose values of *γ* to be as small as possible for the IIT-based model, so that stimulus activity was maintained in a stable high-firing rate attractor state (in line with predictions from the COGITATE consortium) whilst still generating the dip in activity after stimulus-offset that is known to occur empirically, resulting in *γ* = 0.25.

## Supplementary figures

**Figure S1.**
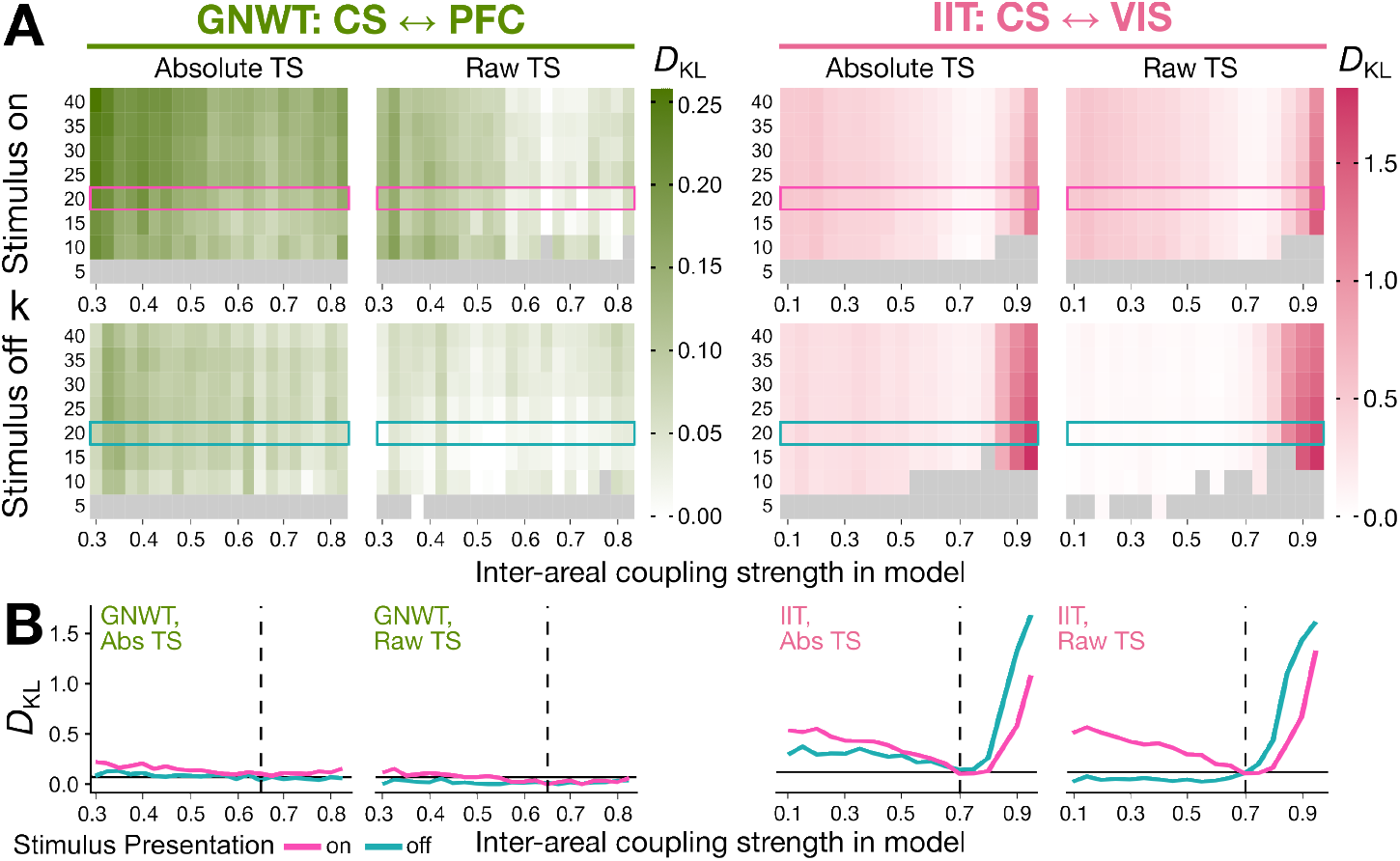
Sweeping model parameters to minimize the *D*_KL_ points to *k* = 20, *W*_GNWT_ = 0.65, and *W*_IIT_ = 0.7 when the added noise *σ*=1. **A**. First, we examined the K-L divergence in timing of the maximum squared Euclidean barycenter between the empirically observed MEG data (*P*) and simulated data for each model (*Q*). For the purpose of examining model parameters, we focus here on the largest added zero-mean noise context (*σ*=1). The *k*-NN estimated K-L divergence is plotted as a heatmap for each inter-areal coupling parameter level (*x*-axis) and each value of *k* nearest neighbors for the estimator (*y*-axis) per model and stimulus presentation period. The row corresponding to *k*=20 is highlighted for all plots, as this is the smallest value of *k* that yielded all finite K-L estimates in all evaluated settings. **B**. The K-L divergence estimated with *k*=20 is plotted for each model and stimulus presentation period, with the inter-areal coupling parameter level on the *x*-axis. For each plot, the dashed vertical line corresponds to the coupling parameter level that minimized the average K-L divergence between stimulus onset (teal) and offset (pink). The resulting minimum average K-L divergence is plotted for each model as a solid horizontal black line. For GNWT, this corresponds to *W*_GNWT_ = 0.65 (with an average *D*_KL_=0.07 nats in the absolute-value time series setting) and *W*_IIT_ = 0.7 (with an average *D*_KL_=0.12 nats in the absolute-value time series setting).

**Figure S2.**
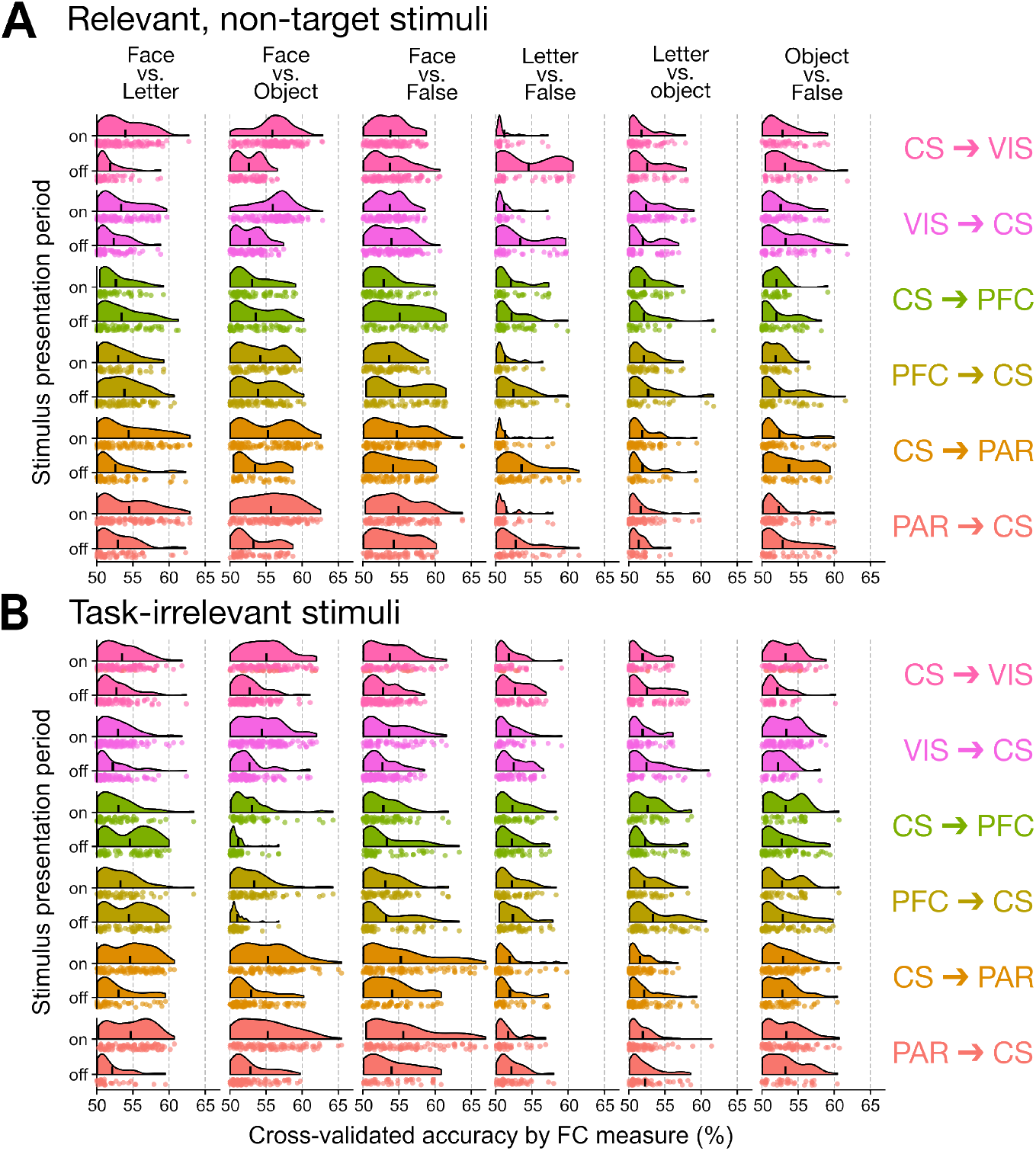
Many FC measures distinguish stimuli with ≥ 60% accuracy across region–region pairs, particularly for face stimuli. **A**. Each raincloud plot shows the distribution of classification accuracy values for each FC measure in the corresponding stimulus pair (column) and region–region pair (row), partitioned by relevant non-target stimulus presentation period (‘on’ vs. ‘off’). Each dot represents one FC measure, and the vertical lines represent the mean cross-validated accuracy across all FC measures in the corresponding distribution. The *x*-axis was truncated at 50% accuracy, such that FC measures yielding lower than 50% accuracy are not shown here. **B**. Same as in **A**, for task-irrelevant stimuli.

**Figure S3.**
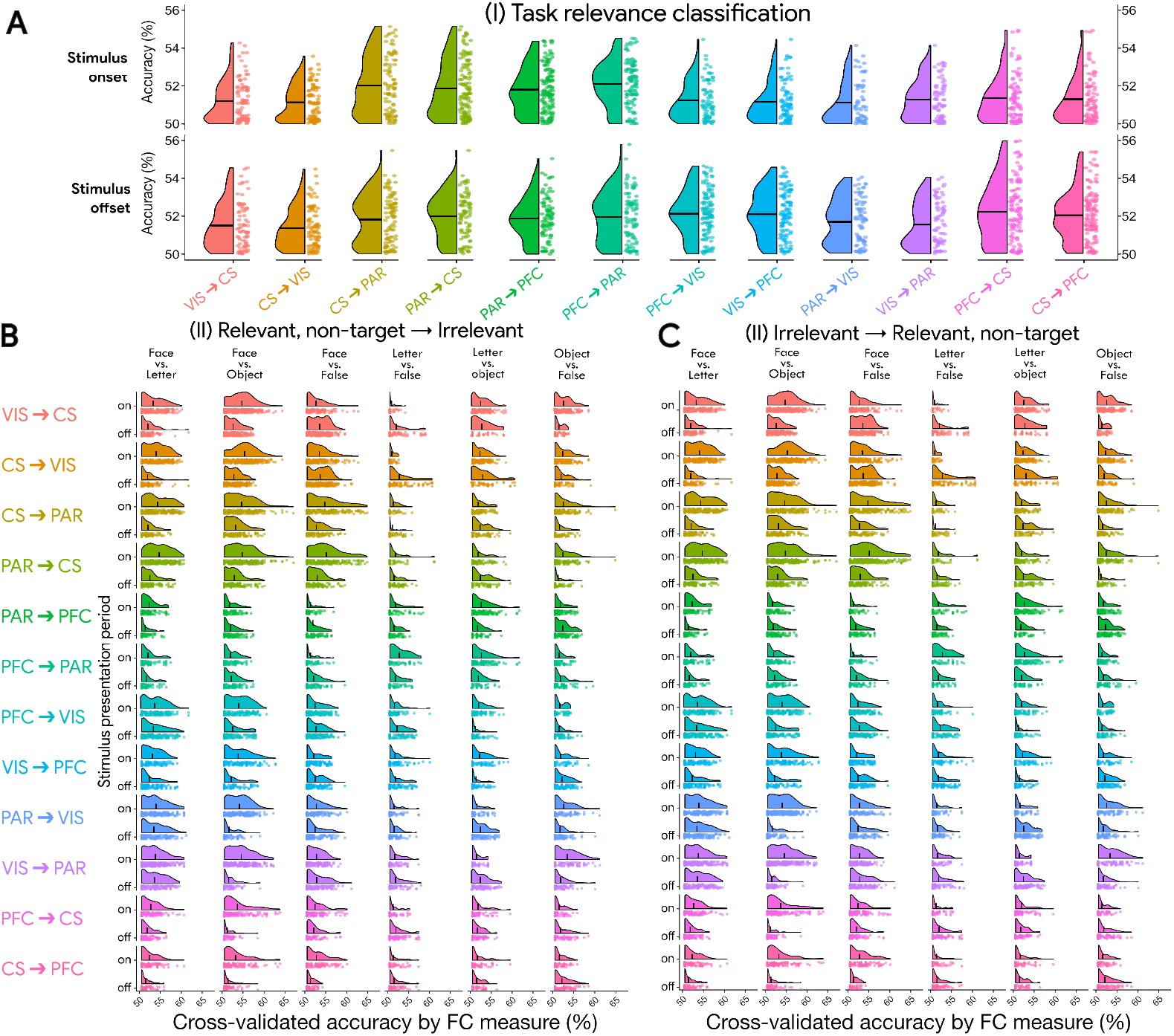
Domain-independent task relevance is not well classified across FC measures and region–region pairs, while cross-task transfer offers comparable performance distributions to the primary stimulus-type classification analysis. **A**. In the first of two supplementary classification aims, we examined how accurately each FC measure could distinguish between ‘Relevant, non-target’ versus ‘Irrelevant’ stimuli in a domain-independent manner. Each raincloud plot shows the distribution of classification accuracy values for each FC measure in the corresponding region–region pair, partitioned by stimulus onset or stimulus offset period. For visualization purposes, the *y*-axis was truncated at 50% accuracy for both stimulus presentation periods, such that FC measures yielding lower than 50% accuracy on average are not shown here. **B**. In the second of two supplementary classification aims, we examined how accurately each FC measure could generalize when trained on a pair of stimuli presented in the context of one relevance type (e.g., ‘Relevant non-target’) and tested on the other relevance type (e.g., ‘Irrelevant’). Here, the distributions show classification accuracy values for each FC measure per region–region pair, trained in the ‘Relevant, non-target’ setting and tested in the ‘Irrelevant’ setting. For visualization purposes, the *x*-axis was truncated at 50% accuracy for both stimulus presentation periods, such that FC measures yielding lower than 50% accuracy on average are not shown here. **C**. The same classification results are plotted as in **B**, here showing the results when trained in the ‘Irrelevant’ setting and tested in the ‘Relevant, non-target’ setting.

**Figure S4.**
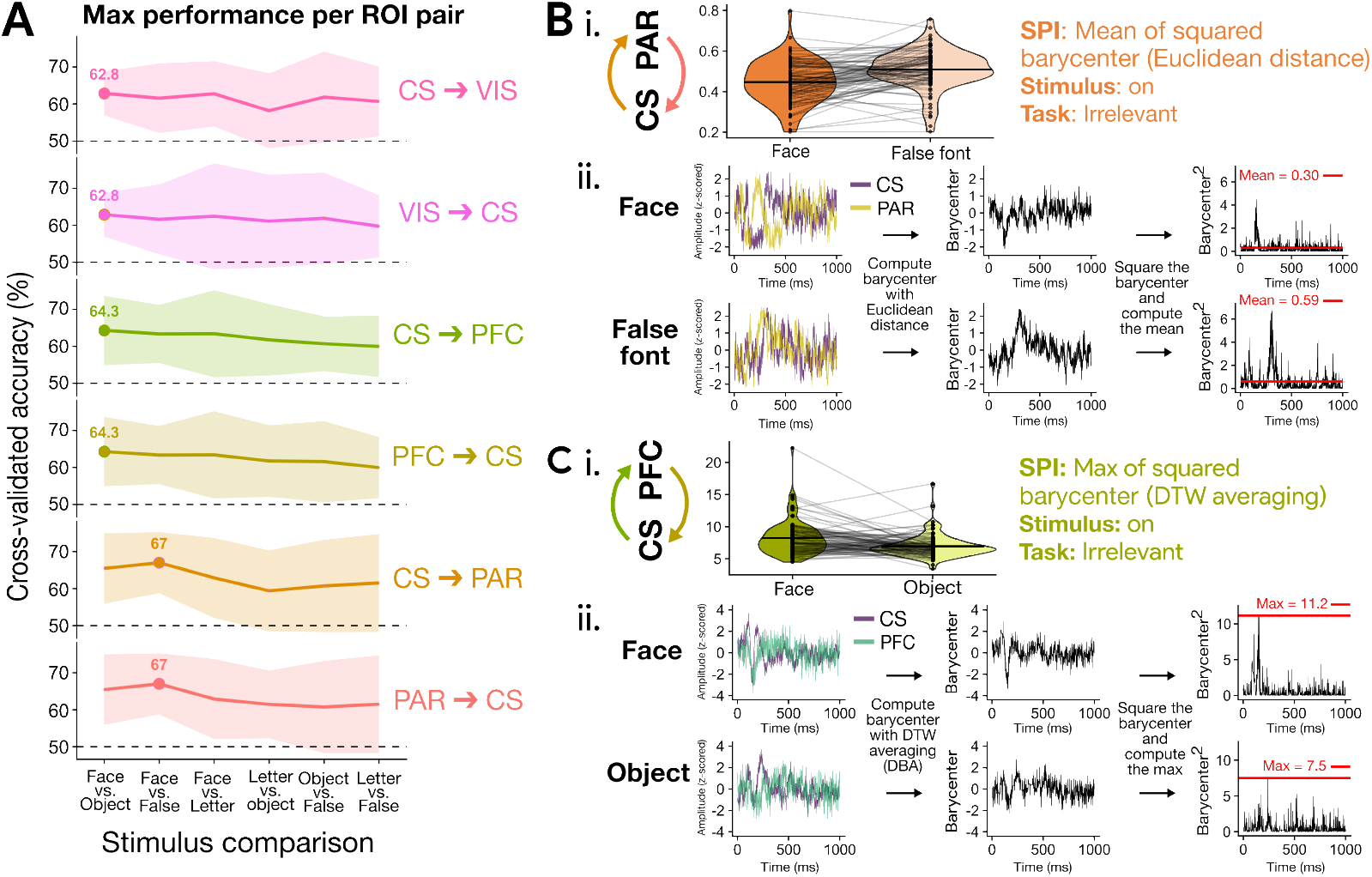
Examining the top-performing FC measure per stimulus comparison highlights strong performance of barycenter statistics. **A**. For each region–region pair (rows) and stimulus comparison (columns), we plot the mean cross-validated classification accuracy for the top-performing FC measure (lines), *±* 1 SD (shaded ribbons) across test folds, out of all 246 FC measures, relevance conditions (‘Relevant non-target’ or ‘Irrelevant’), and stimulus presentation periods (‘on’ or ‘off’). The peak classification accuracy is annotated per region–region pair. **B**. (i) The top-performing FC measure for CS ←→ PFC—the maximum of the squared barycenter with DTW averaging—is schematically depicted. The violin plots show the FC measure value distributions for task-irrelevant ‘face’ versus ‘object’ trials during stimulus onset across all N=94 participants. (ii) Below, line plots schematically depict the computation of this FC measure from ‘face’ or ‘object’ trial-averaged MEG time series in an example participant. **C**. (i) as in **B**(i), the top-performing FC measure for CS ←→ PAR—the mean of the squared barycenter with Euclidean geometry—is schematically depicted. The violin plots show the FC measure value distributions for task-irrelevant ‘face’ versus ‘false font’ trials during stimulus onset across all N=94 participants. (ii) Below, line plots schematically depict the computation of this FC measure from ‘face’ or ‘false font’ trial-averaged MEG time series in an example participant (n.b., a different participant than in (i)).

**Figure S5.**
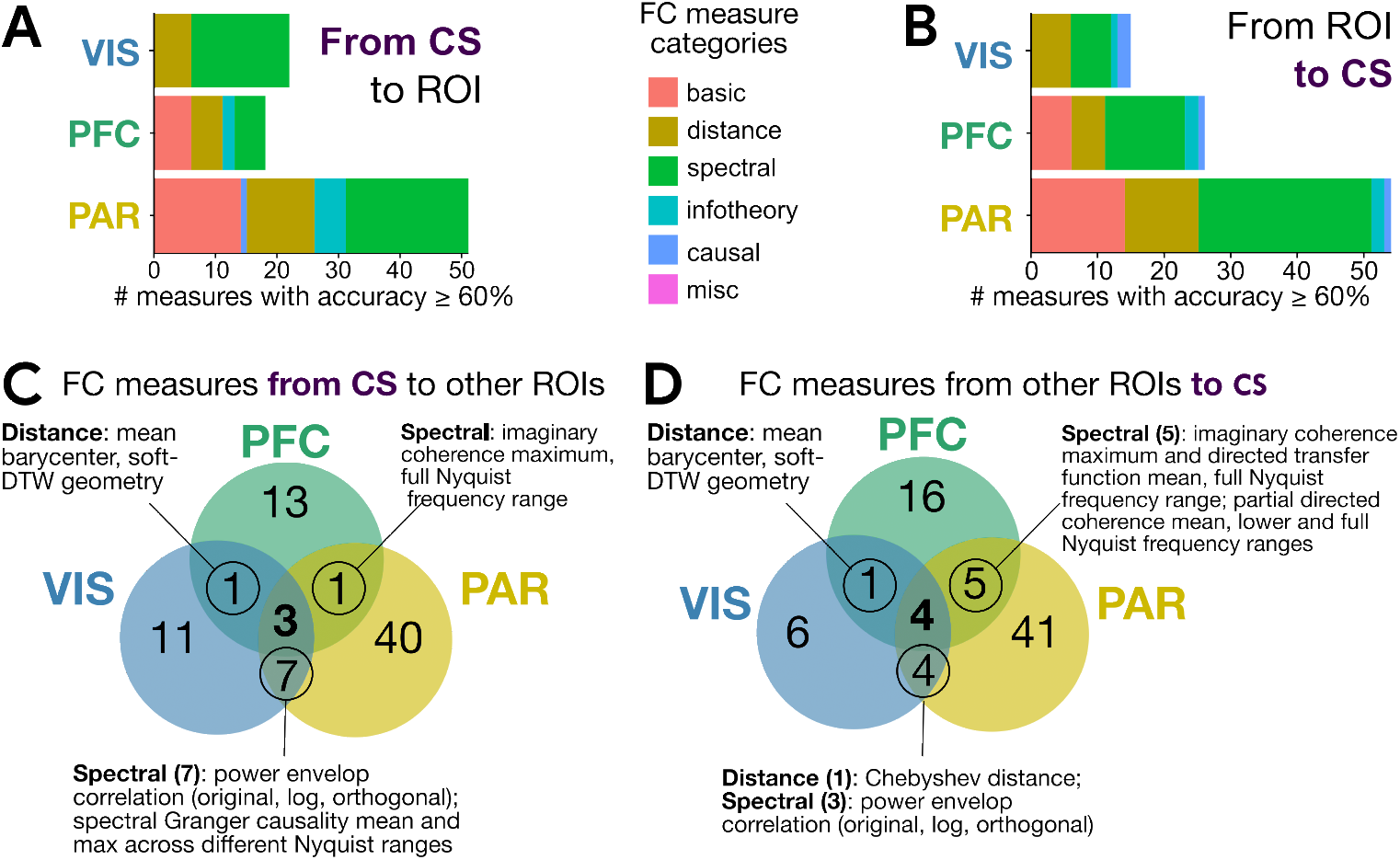
Detailed breakdown of the top-performing FC measures shared between two functional pathways. **A, B**. For the FC measures tabulated in Fig. 4A and Fig. 4B, here we show the number of FC measures by literature category per region as the source (**A**) and target (**B**). **C**. The same Venn diagram from Fig. 4A is reproduced, with detailed annotation of the FC measures comprising the intersection between two functional pathways projecting **from** CS to VIS, PFC, or PAR. **D**. The same Venn diagram from Fig. 4B is reproduced, with detailed annotation of the FC measures comprising the intersection between two functional pathways projecting **to** CS from VIS, PFC, or PAR.

**Figure S6.**
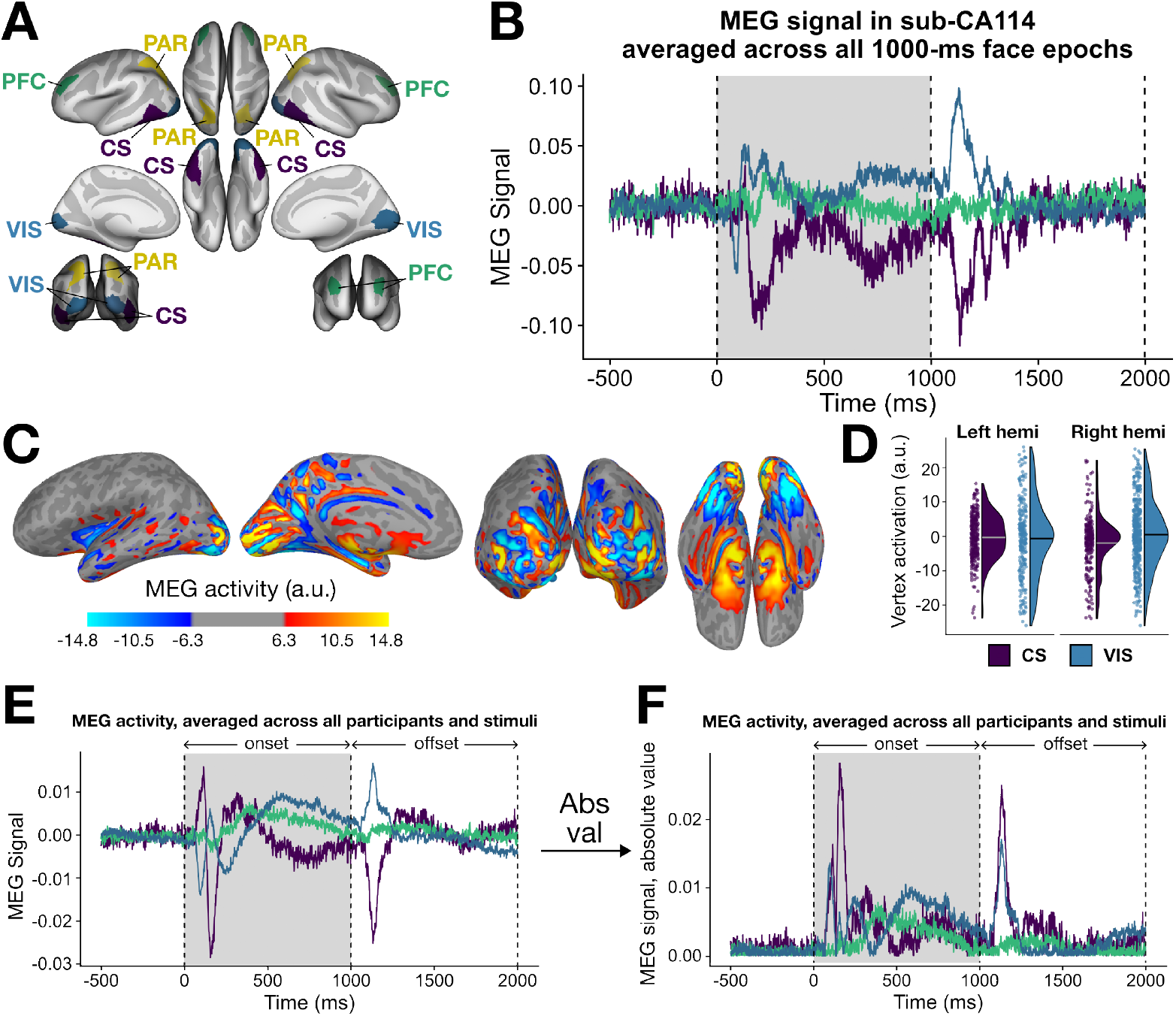
Analyzing regional polarity in MEG signal supports computing the absolute value of MEG amplitude. **A**. The four regions of interest examined in this study are shown on the cortical surface in `fsaverage` space. **B**. The average MEG signal is shown for one participant (sub-CA114), with activity averaged across all epochs per region. The regions are colored as in **A**. Stimulus onset is denoted with a shaded gray rectangle (occurring between 0 to 1000 ms), followed by the 1000-ms offset period. **C**. The raw MEG signal is plotted at the point of peak brain-wide intensity, corresponding to 153 ms post-onset. Larger activity magnitudes indicate greater electrical current, arising primarily from postsynaptic activity in pyramidal neurons in the cortex, with the sign indicating the orientation relative to the MEG sensors. **D**. Vertex-wise activation values are summarized in raincloud plots for the CS and VIS regions in the left and right hemispheres, respectively. Horizontal bars in each half-violin represent the mean activation value in artificial units (a.u.). **E**. The average MEG signal is shown, computed across all participants (N=94) and all epoch types, per brain region. The regions are colored as in **A**. Stimulus onset is denoted with a shaded gray rectangle (occurring between 0 to 1000ms), followed by the 1000-ms offset period. **F**. The absolute value is obtained from the average MEG time series shown in **E**.

**Figure S7.**
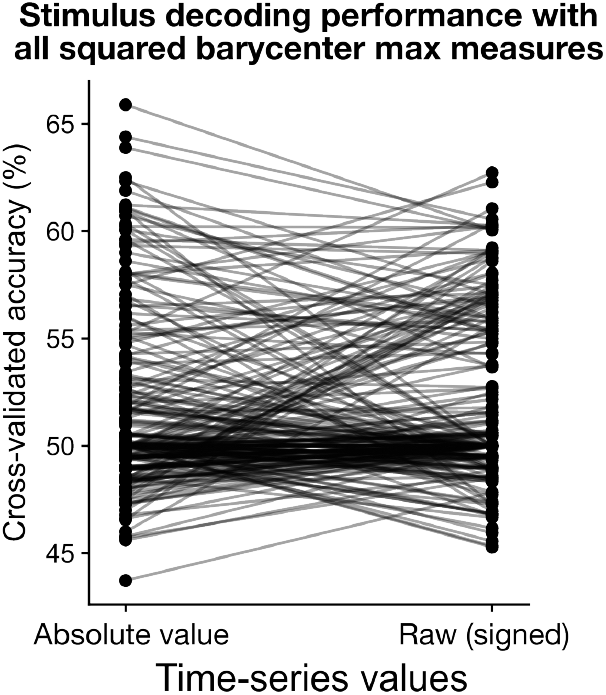
Stimulus decoding performance with the maximum of the squared barycenter (across all four evaluated distance metric geometries) does not systematically vary between the absolute value versus raw (signed) empirical MEG time series.

**Figure S8.**
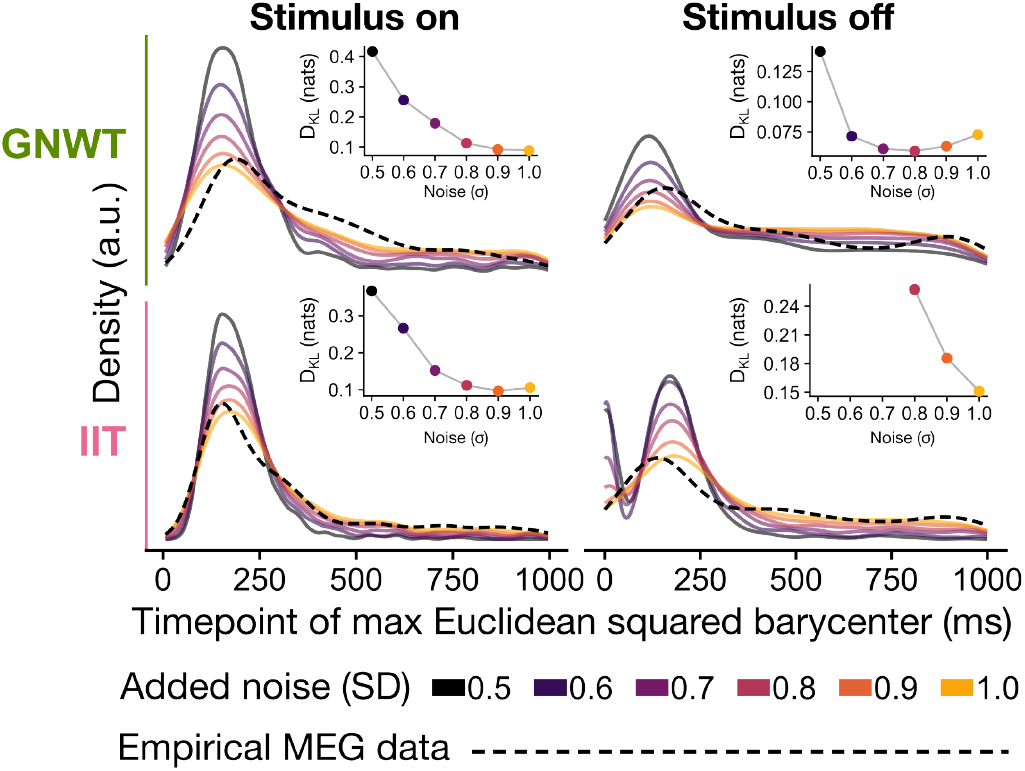
K-L divergence patterns are robust across added noise levels for simulated time series between GNWT and IIT. In each stimulus presentation period (‘on’ vs. ‘off’) and model context (‘GNWT’ vs. ‘IIT’), the distribution of timepoints for the maximum squared Euclidean barycenter is plotted for the empirical MEG data (thicker dashed black line) as well as for model-simulated data with noise thresholds ranging from 0.5 ≤ *σ* ≤ 1.0. The inset in each graph depicts the K-L divergence between the empirical versus model-simulated time series at the corresponding noise level.

**Figure S9.**
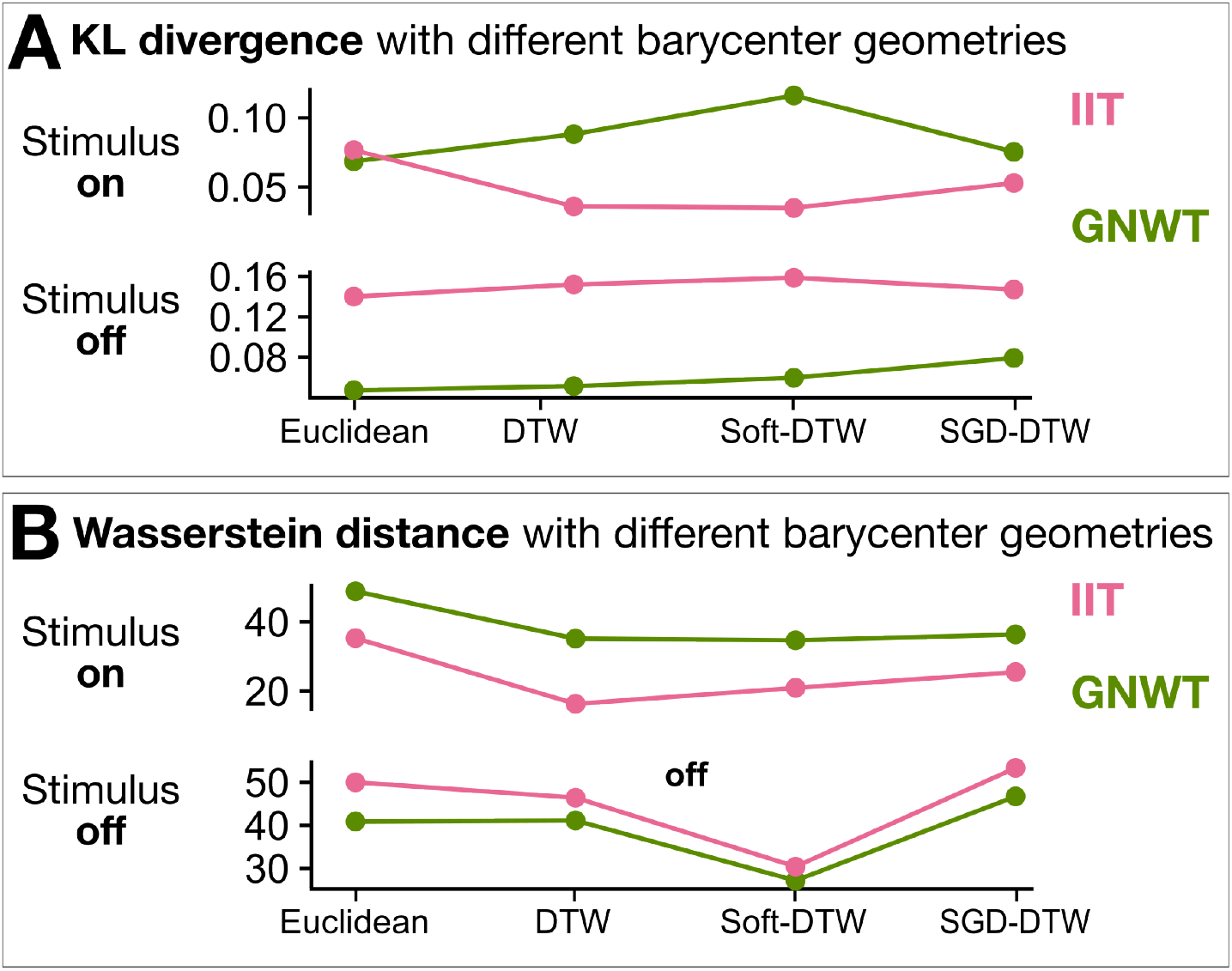
K-L divergence and Wasserstein distance between empirical vs. model-based maximum squared barycenter timepoints show consistent trends across different Euclidean and non-Euclidean barycenter geometries.

**Figure S10.**
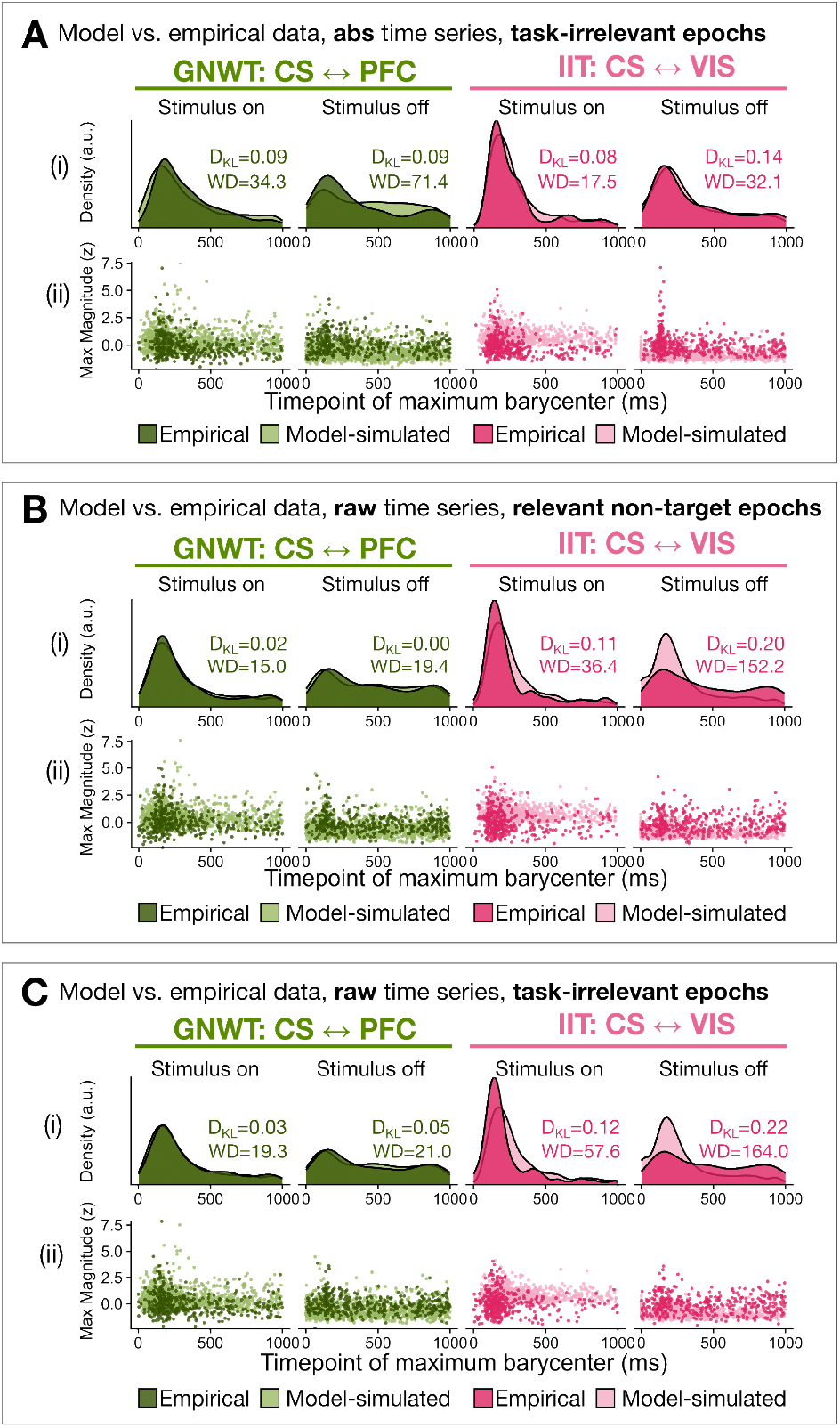
Timing of the maximum squared Euclidean center is better approximated by the GNWT-based than IIT-based model across stimulus relevance settings, with either raw or absolute value time series. In each setting, we compare the distribution of timepoints corresponding to the maximum squared Euclidean barycenter for the empirical data (darker shading) versus model-simulated time series (lighter shading) for GNWT in green, corresponding to CS ↔ PFC, and for IIT in pink, corresponding to CS ↔ VIS in pink. The kernel-estimated densities are shown in each panel in (i), with the K-L divergence (*D*_*KL*_) and Wasserstein distance (WD) given for each empirical vs. model-simulated comparison. The timepoints and corresponding maximum barycenter magnitudes are shown in each (ii) panel. Here, we compare (**A**) absolute time series in task-irrelevant epochs; (**B**) raw time series in relevant non-target epochs; and (**C**) raw time series in task-irrelevant epochs.

## Notes

https://github.com/anniegbryant/MEG_functional_connectivity

https://doi.org/10.17617/1.WQA3-WK71

### Competing Interest Statement

The authors have declared no competing interest.

### Summary of Updates

Introduction revised to more thoroughly introduce the barycenter as a concept. Methods have been revised to more thoroughly explain region of interest selection and to include area under the curve analysis, RBF kernel in addition to linear kernel for SVM robustness analysis, kNN-estimated Kullback-Leibler divergence, and more rigorous parameter sweeps for the neural mass models. Results have been revised to visually refine figures and to reflect the new KL divergence estimates and the new parameter configurations for the neural mass models. Discussion has been revised to contextually embed the magnitude of classification performances obtained here and to flow in a more organized manner.

